# Winter warming shapes gut microbiome composition and directional dysbiosis in a temperate lizard

**DOI:** 10.64898/2026.02.20.706997

**Authors:** Miary Raselimanana, Amy Ellison, Alberto H. Orta, John W. Wilkinson, Wolfgang Wüster, Kirsty J. MacLeod

## Abstract

Environmental temperature shapes ectotherm gut microbiomes through direct effects on community structure and indirect effects mediated by host physiology. Due to climate change, winter temperatures are rising, in some regions faster than summer temperatures. However, warming effects on the microbiome during overwintering remain poorly understood compared to the active season, limiting predictions of host-microbiome responses across the annual cycle. We experimentally tested how winter warming influences gut microbiome diversity and composition in the common wall lizard (*Podarcis muralis*). Thirty-nine lizards were overwintered under cold (4±1°C), mild (8±1°C), or fluctuating (5 days cold, 2 days mild) temperatures, and 80 faecal samples were collected post-emergence for 16S rRNA gene sequencing. Winter warming did not alter alpha diversity but induced consistent shifts in community composition relative to the cold treatment, with distinct responses to constant versus fluctuating warming. Constant mild temperatures enriched fewer taxa, including putatively opportunistic or pathogenic genera, potentially signalling microbiome imbalance. In contrast, fluctuating warming and the baseline cold treatment preserved a broader suite of fermentative bacteria, likely supporting more stable gut homeostasis. Both warming treatments increased directional dysbiosis relative to the cold treatment, though without increasing interindividual variability, indicating structured reassembly rather than stochastic change. Our findings suggest that winter warming can subtly negatively affect gut microbiomes and host health, while fluctuating temperatures may buffer negative effects. We show that overwintering is an important, yet overlooked, period through which climate warming can shape host-microbiome dynamics.

## 1. Introduction

Gut microbial communities are largely shaped by the internal environment created by their hosts (Jandhyala et al., 2015; Tremaroli and Bäckhed, 2012), and in turn influence host physiology, development and fitness (e.g., McFall-Ngai et al., 2013; Zhang et al., 2019). In ectothermic hosts, whose physiological processes depend strongly on ambient temperature, gut microbiome composition is additionally sensitive to external thermal conditions (e.g., Kohl and Yahn, 2016; Moeller et al., 2020; Sepulveda and Moeller, 2020). Temperature shifts can alter the relative abundance of key microbial taxa, with warming often reducing beneficial, disease-resistant bacteria while promoting relatively thermotolerant pathogenic or opportunistic taxa, defined here as taxa that can cause disease under certain conditions (Almblad et al., 2021; Brewer et al., 2021; Fontaine et al., 2018). These microbial changes can cascade to affect host physiology, immune function, behaviour, and overall fitness (Comizzoli et al., 2021; Greenspan et al., 2020; Hanning and Diaz-Sanchez, 2015; Hector et al., 2022; McFall-Ngai et al., 2013; Tremaroli and Bäckhed, 2012). Because gut microbiota mediate host responses to environmental stressors (Rocca et al., 2019), including thermal variation, they play a central role in determining organismal resilience under climate change (Bestion et al., 2017; Ferguson et al., 2018a; Hylander and Repasky, 2019; Olsson et al., 2025).

In squamate reptiles, gut microbiota respond dynamically to environmental conditions (Hoffbeck et al., 2023; Kohl et al., 2017), with temperature emerging as a particularly strong driver of community structure. Across species, warming has been repeatedly associated with reduced microbiome stability, losses of core taxa, and increases in opportunistic or potentially pathogenic bacteria (e.g., Moeller et al., 2020; Shi et al., 2025; Zhang et al., 2022). For example, warming reduced *Bacillota* abundance, decreased within-individual microbiome stability, and increased predicted pathogens in western fence lizards (*Sceloporus occidentalis*) (Moeller et al., 2020). Even modest warming (2-3 °C) caused substantial and persistent loss of core microbiota diversity in common lizards (*Zootoca vivipara*) (Bestion et al., 2017).

These shifts may arise because many bacterial lineages are highly sensitive to temperature and can directly perceive small thermal changes, adjusting growth rates or virulence accordingly (Kim et al., 2019; Roncarati et al., 2025). The loss of core taxa can in turn facilitate pathogen colonisation and impair nutrient absorption (Kohl and Yahn, 2016). Importantly, the observed patterns reflect not only compositional change but a reduction in host regulation of the microbiome, often accompanied by increased among-individual variability in community structure. This is consistent with the Anna Karenina Principle (AKP), which predicts that environmental stress destabilises host-microbe interactions, leading to increased community dispersion and/or interindividual host variability in microbiome structure (Greenspan et al., 2020; Lesser et al., 2016; MacLeod et al., 2022; Stothart et al., 2016; Zaneveld et al., 2017). Hereafter, we refer to this pattern of instability and increased dispersion/variability as dysbiosis (Arnault et al., 2023; Greenspan et al., 2020; Wei et al., 2021). However, existing studies have focused almost exclusively on active season warming, when hosts are feeding and metabolically active, and thus may not capture microbiome responses under the distinct physiological state associated with overwintering. Whether winter warming constitutes a comparable environmental stressor capable of inducing AKP-type destabilisation therefore remains unclear (MacLeod et al., 2022).

Winter represents a fundamentally different physiological context for ectotherms. Many temperate species enter complete or partial dormancy to conserve energy during periods of low resource availability (Blix, 2016; Wilsterman et al., 2021), relying on stored reserves to survive prolonged fasting (Bonnet et al., 1998; Shine, 2005). Because host physiology during winter differs markedly from the active season, microbiome responses to winter warming may also differ. Importantly, winter temperatures have increased at a faster rate than summer, with more frequent warm spells and greater thermal variability (Intergovernmental Panel on Climate Change, 2014; Vose et al., 2005), disrupting overwintering ectotherms (Williams et al., 2015), including squamate reptiles (Moss and MacLeod, 2022; Raselimanana et al., 2025; Turner and Maclean, 2022). Moreover, seasonal studies indicate that gut microbiomes of overwintering squamates naturally fluctuate across seasons, suggesting sensitivity to winter physiological states (Zhu et al., 2024). Yet, the consequences of winter-warming – and of different winter warming patterns – for gut microbiome diversity, composition, and stability remain poorly understood. Given predictions of increasing mean winter temperatures and thermal variability (Lee et al., 2023), it is therefore essential to test whether different overwintering temperature regimes (e.g., constant mild warming versus fluctuating temperatures) lead to divergent microbiome responses.

To address these gaps, we experimentally manipulated overwintering temperatures (cold, mild, fluctuating) in a temperate-zone squamate (the common wall lizard, *Podarcis muralis*; Laurenti, 1768) to examine how winter thermal conditions shape gut microbiome diversity and community composition, and whether there is evidence for microbiome dysbiosis under warming conditions. We predicted that: (i) alpha diversity would decrease under mild and fluctuating winter temperature regimes relative to cold conditions, reflecting the sensitivity of gut microbial diversity to environmental stressors (Rocca et al., 2019); and (ii) gut microbiome composition (beta diversity) would shift in response to winter warming, increasing community dispersion and interindividual variability, consistent with greater host-associated microbiome dysbiosis. We further investigated potential sex-specific effects, as sex can influence host-associated microbial communities in ectotherms (Bunker et al., 2022; Martin et al., 2010; Zhang et al., 2022a; Zhou et al., 2020). Finally, we examined temporal dynamics following emergence from overwintering, predicting that winter temperature-induced microbial differences would persist into the early active season. To our knowledge, this study provides the first experimental assessment of how contrasting winter-warming regimes shape gut microbiome diversity and composition in overwintering reptiles. By exploring winter temperature as a potentially important but understudied determinant of host-microbiota interactions, we aim to advance predictions of ectotherm resilience to ongoing climate warming (Altizer et al., 2013; Bestion et al., 2017).

## 2. Materials and methods

### 2.1 Study species

The common wall lizard (*Podarcis muralis*) is a small lacertid lizard widely distributed across Europe and introduced in several regions, including the United Kingdom (Kolenda et al., 2020; Michaelides et al., 2013; Michaelides et al., 2015; Oskyrko et al., 2020; Schulte et al., 2012). UK populations occur at the northern edge of the species’ range and likely rely on winter dormancy, as in other cool climates, because temperatures frequently fall below activity thresholds, despite occasional activity during warm spells (Burke and Ner, 2005; Foa et al., 1992; Sakelarieva et al., 2023).

Lizards were captured from an introduced population in Bournemouth, Dorset, England (50.72°N, -1.82°W, 10-20 m a.s.l.), UK. Thirty-nine adults (23 males, 16 females) were captured during early autumn 2022 using a combination of hand capture and lassoing, and transported to Bangor University, Wales, within 72 hours.

### 2.2 Overwintering experiment

Lizards were overwintered under one of three experimental thermal treatments representing a gradient from cold to mild winter conditions: control cold winter (4±1°C), constant mild winter (8±1°C), and fluctuating winter temperatures (alternating 5 days at 4±1°C and 2 days at 8±1°C). These thermal regimes reflect ecologically relevant temperatures near the boundary between physiological torpor and intermittent winter activity (Barroso, 2023; While et al., 2015), with the control treatment approximating historical mean winter air temperature in southern UK (3.73°C, 1991-2020, metoffice.gov.uk). The experiment ran for 104 days (Figure 1).

**Figure 1:**
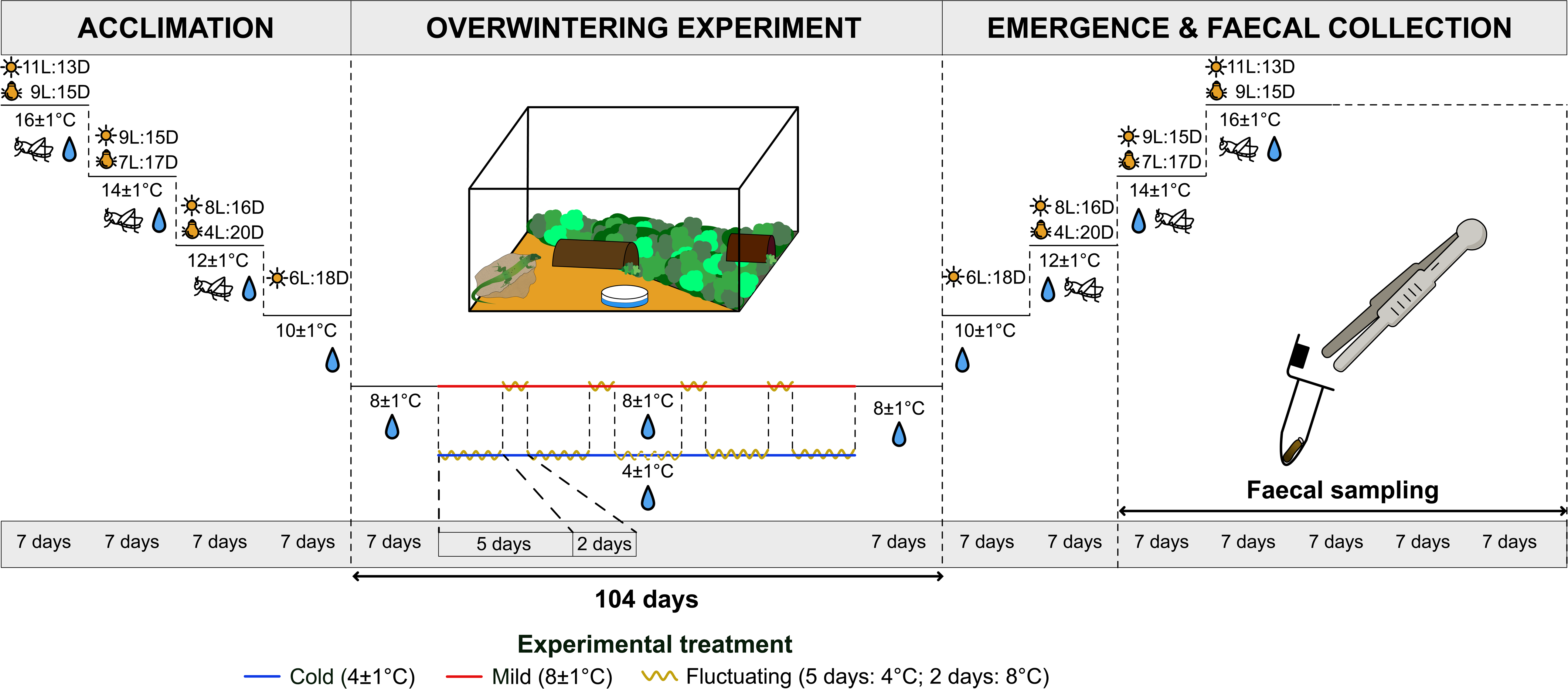
Diagram illustrating the experimental design, comprising the acclimation, overwintering, and post-overwintering periods for common wall lizards (*Podarcis muralis*), with durations indicated in days. Photoperiod from the main room light (sun icon), and UV basking light (bulb icon) is shown with light: dark (L:D) cycles (hours) indicated where they change. Temperature was gradually adjusted pre- and post-overwintering and over 7 days at the start and towards the end of overwintering, then maintained for the cold (4±1°C; blue line) and mild treatments (8±1°C; red line) and changed for the fluctuating treatment (5 days at 4°C and 2 days at 8°C; yellow zigzag line). Resource availability is shown with a cricket icon for food and a water droplet icon for water.

Individuals were marked with non-toxic paint on the dorsal or lateral surface for identification and re-marked as needed. Prior to overwintering, lizards underwent a ∼30-day acclimation period during which temperature and photoperiod were gradually reduced to induce torpor as described in Raselimanana et al. (2025). At the end of the experiment, pre-hibernation conditions were gradually restored (Figure 1). Lizards from the same treatment were then randomly assigned to communal plastic terraria in groups of 3-5, with no more than two males per enclosure to minimise aggression (Sacchi et al., 2009). Feeding resumed five days after emergence.

We measured snout-vent length (SVL) and body mass of the lizards after emergence to assess host body condition. Body condition was quantified as the residuals of the linear regression of log_10_-transformed body mass on log_10_-transformed SVL, calculated separately for males and females (Amer et al., 2024; Perry et al., 2024; Schulte-Hostedde et al., 2005).

### 2.3 Faecal sample collection

Faecal sampling began one week after feeding resumed (ensuring the collection of the first post-hibernation faeces) and was continued for five weeks (Figure 1). On scheduled sampling days, lizards were temporarily placed in individual plastic containers lined with clean paper substrate in the late afternoon (16:00-18:00). Faeces were collected the following morning using either stainless-steel forceps sterilised with 95% ethanol between samples or single use sterilised plastic scoops. Gloves were changed between each collection to minimise the risk of cross-contamination, after which lizards were returned to their original communal terraria. When defecation occurred during relocation or within the communal terrarium, only freshly deposited faeces observed less than 1h after defecation were collected. Small amounts of soil substrate were also sampled randomly from terraria as environmental controls to account for bacterial communities present in the substrate. All faecal (*n* = 80) and control substrate samples (*n* = 4) were immediately frozen at -80°C following collection.

Sampling was conducted approximately three times per week, yielding up to ten sampling opportunities per individual across the five-week period. Because defecation frequency varied among individuals, not all lizards contributed samples at each time point, and some individuals provided multiple samples. This imbalance was accounted for in downstream statistical analyses by including individual identity as a random effect. Sampling effort across treatments, sexes and time points is summarised in Table S1. Seven samples lacked a clearly recorded collection date and were therefore excluded from temporal analyses. Additionally, 14 samples collected ≤1h after defecation in the communal terraria could only be assigned to the terrarium (and thus treatment level) rather than to individual lizards. To account for uneven sampling effort among collection days and to facilitate analyses of temporal dynamics, samples were aggregated to weekly time points. To further increase sample sizes at later stages of faecal collection, all samples collected from week 3 onwards were pooled into a single category. Consequently, temporal analyses were conducted using three time points: week 1, week 2, and weeks 3+.

### 2.4 DNA extraction, 16S rRNA amplification and sequencing

DNA was extracted from 80 faecal samples and 4 substrate controls using the DNeasy PowerSoil Pro Kit (QIAGEN, Venlo, The Netherlands) following a validation comparison with the QIAamp Fast DNA stool mini kit (QIAGEN, Venlo, The Netherlands) (see Supplementary Method). We also conducted six blank extractions (1 blank per batch of sample) to monitor contamination and correct for contaminants in DNA extraction kits (Salter et al., 2014). Samples were randomised across extraction batches. Extracted DNA was stored at −20°C until Polymerase Chain Reaction (PCR) amplification and library preparation. The V4 region of the 16S rRNA gene (∼253 bp) was amplified for Illumina sequencing (full molecular protocols are available in Supplementary Method). We sequenced bacterial 16S rRNA amplicons from 80 samples, one substrate sample, six extraction blanks, six PCR negative controls, and one substrate-extraction blank.

### 2.5 Bioinformatics

Raw reads were demultiplexed with *cutadapt* v5.1 (Martin, 2011) and processed using the DADA2 pipeline (Callahan et al., 2016). Sequences outside the expected V4 amplicon length (200-300 bp) were discarded. Taxonomy was assigned using DADA2’s naïve Bayesian classifier (Callahan et al., 2016) trained on the SILVA 16S reference database (v138.2; Quast et al., 2013). Low-abundance amplicon sequence variants (ASVs; ≤ 10 total reads), ASVs lacking phylum-level classification, and non-bacterial sequences (chloroplasts, mitochondria) were removed. Contaminant ASVs were identified as those detected in extraction blanks, PCR negatives, or substrate controls, and subsequently removed. Detailed bioinformatic parameters are provided in the Supplementary Methods.

We rarefied ASV tables to 8,000 sequences before comparisons of alpha and beta diversity metrics, following recommendations for depthl1lnormalised diversity estimation (Schloss, 2024). Rarefaction was used only for diversity-based analyses. All other analyses were conducted on non-rarefied data. After quality control and removal of non-bacterial sequences, our sequencing efforts yielded a total of 2.98 million sequences (mean ± SE: 37,196 ± 1979 per sample). These sequences were grouped into 615 ASVs across 80 samples, with rarefaction retaining 503 ASVs across 78 samples.

### 2.6 Statistical analysis

#### 2.6.1 Gut microbiome alpha diversity analysis

We quantified gut microbiome alpha diversity using three complementary metrics: richness (total number of ASVs), the Shannon diversity index (Shannon, 1948), and Faith’s phylogenetic diversity (PD) (Faith, 1992). The Shannon index considers both richness and evenness, and Faith’s PD accounts for evolutionary relationships among taxa based on total branch length in the ASV phylogeny. Observed ASVs and the Shannon index were calculated using *phyloseq*, and Faith’s PD using *picante* v1.8.2 (Kembel et al., 2010).

To test whether overwintering temperature affected gut microbiome diversity, we fitted separate linear mixed-effects models (LMMs) for each alpha diversity metric using *lme4* v1.1-37 (Bates et al., 2015). Fixed effects included temperature treatment (cold, mild, fluctuating), sampling time points (week 1, week 2, week 3+), and sex. Individual ID and terrarium ID were included as random intercepts to account for repeated measures and potential grouping effects. Random effects were retained only when they explained non-negligible variance and did not result in singular model fits. Interaction terms (treatment×sex, treatment×time) were evaluated but not retained in the final models due to limited sample sizes and lack of improvement in overall model performance (ΔAIC ≥ 2). Model assumptions were assessed graphically and using diagnostic tests from *DHARMa* v0.4.7 (Hartig, 2024). Because host body condition has been shown to predict microbiome diversity and composition (e.g., Chun et al., 2020; Hoffbeck et al., 2024; Ruiz-Rodríguez et al., 2009; Slevin et al., 2025), body condition at emergence was initially considered as a covariate. However, body condition differed significantly among overwintering treatments (*F*_2,62_ = 13.76, *p* < 0.001), indicating that it was not independent of the experimental manipulation. To avoid collinearity, body condition was therefore excluded from the final models, and associations between condition and microbiome responses, including alpha diversity and dysbiosis, were instead examined using correlation analyses (see section 2.6.4).

#### 2.6.2 Compositional (beta diversity) analysis

We quantified gut microbiome compositional differences using Bray-Curtis dissimilarity (Bray and Curtis, 1957) and unweighted UniFrac distances (Lozupone and Knight, 2005), calculated in *phyloseq*. Bray-Curtis captures abundance-based community dissimilarity, whereas unweighted UniFrac incorporates phylogenetic relationships based on the presence-absence of ASVs. Differences in community structure were visualised with principal coordinates analysis (PCoA) using *ape* v5.8-1 (Paradis and Schliep, 2019).

To test whether overwintering temperature treatment affected gut microbiome composition, we conducted permutational multivariate analysis of variance (PERMANOVA) using *vegan::adonis2* (Oksanen et al., 2025) with treatment (cold, mild, fluctuating), sampling time point (week 1, week 2, week 3+), and sex as fixed effects. LizardID was included as a blocking factor to control for repeated sampling of individuals. Separate PERMANOVA models were run for each distance metric. Non-significant interaction terms (treatment×time, treatment×sex) were removed to facilitate interpretation of main effects (full model output in Table S3).

We also performed differential abundance analysis using *DESeq2* v1.46.0 (Love et al., 2014) on non-rarefied ASV count data to identify bacterial taxa associated with treatment, time, and sex. Statistical analyses were conducted at the ASV level, while taxonomic interpretation of significant results was performed at the genus level. For visualisation, significant ASVs were further aggregated at the order level to improve readability, with genus-level plots provided in Supplementary Figure S1. To ensure robustness of taxonomic inference, we retained for each contrast only genera supported by at least two ASVs (adjusted *p* < 0.05) or by a single significant ASV that was highly abundant in the community (i.e. relative abundance ≥ 10%).

#### 2.6.3 Dysbiosis: community dispersion, directional shift, and interindividual variability

We quantified dysbiosis using three complementary metrics capturing distinct dimensions of microbiome disruption: (i) within-group dispersion (community heterogeneity), (ii) directional deviation from a reference community, and (iii) interindividual variability. Together, these metrics characterise changes in microbiome stability, divergence, and consistency across individuals. First, we assessed whether overwintering temperatures altered microbiome heterogeneity by quantifying multivariate dispersion of beta diversity distances (Bray-Curtis and unweighted UniFrac) using *vegan::betadisper* (Oksanen et al., 2025). Dispersion values (distance to group centroid) were analysed using linear models with treatment, sex, and their interaction as fixed effects.

Second, we quantified directional shifts in microbiome composition by calculating a dysbiosis score for each sample as its Euclidean distance to the centroid of a reference community. The cold treatment was used as a baseline representing typical overwintering conditions (see section 2.2). Distances were calculated using the *dysbiosisR* package v1.0.4 (Shetty, 2022) and analysed with linear models including treatment, sex, and their interaction as fixed effects. This metric captures the magnitude of community deviation independently of differences in dispersion.

For both dispersion and directional dysbiosis metrics, we also calculated area-under-the-curve (AUC) values using *pROC* v1.19.0.1 (Robin et al., 2011) as a descriptive measure of how well each metric distinguished treatments. A value of ≥ 0.80 indicates good discrimination performance.

Finally, interindividual variability was quantified as the mean pairwise distance between each sample and all other samples within the same treatment group (MacLeod et al., 2022) and analysed using mixedl1leffects models. As no treatment effects were detected, these results are presented in the Supplementary Results.

#### 2.6.4 Correlations between host body condition and microbiome diversity and dysbiosis

To address the possibility that gut microbiome responses to winter-warming would be condition-dependent, we hypothesised that individuals in better body condition (body condition index > 0) would show more diverse and less dysbiotic microbiomes. This prediction is consistent with previous studies demonstrating condition-dependent microbiome effects in birds and reptiles (Ruiz-Rodríguez et al., 2009; Slevin et al., 2025; Teyssier et al., 2018). We explored potential condition-dependent effects using separate correlation analyses between body condition and gut microbiome metrics, which allowed us to preserve degrees of freedom in the main models (sections 2.6.1 and 2.6.3). Alpha diversity metrics included observed ASVs, the Shannon diversity index, and Faith’s phylogenetic diversity, whereas dysbiosis metrics included community dispersion, interindividual variability, and directional dysbiosis score (as defined above). Analyses were conducted separately for each sex using Spearman’s rank correlations with Benjamini-Hochberg false discovery rate (FDR) correction (Benjamini and Hochberg, 1995). No significant associations with body condition and any diversity or dysbiosis metric were detected (see Supplementary Results) and are not discussed further.

## 3. Results

### 3.1 Winter warming regimes do not alter gut microbiome alpha diversity

We analysed 80 faecal samples, including 66 samples collected from 24 identified adult common wall lizards (9 females, 15 males) and 14 opportunistic samples collected from communal terraria where individual identity could not be assigned (Table S1). Each individual contributed a maximum of five samples across the five-week post-emergence period, resulting in uneven sampling across sex and time (Table S1). Across all samples, 16 bacterial phyla were detected in the gut, with communities dominated by Bacteroidota (28.7%), Bacillota (25.2%), and Pseudomonadota (12.3%) (Figure S2).

Based on the final supported models, we found no evidence that overwintering temperature treatment (cold, mild, fluctuating) influenced gut microbiome alpha diversity (observed ASVs: *F*_2,54_ = 0.35, *p* = 0.708; Shannon diversity index: *F*_2,52_ = 0.13, *p* = 0.874; Faith’s Phylogenetic Diversity: *F*_2,20.5_ = 0.37, *p* = 0.694). Alpha diversity also did not vary across the sampling time for any metric retained in the final model (Shannon diversity index: *F*_2,52_ = 0.05, *p* = 0.953; Faith’s PD: *F*_2,44.1_ = 1.21, *p* = 0.308), indicating temporal stability in within-sampled diversity.

In contrast, alpha diversity differed between sexes. Females exhibited higher ASV richness and greater phylogenetic diversity than males (Observed ASVs: *F*_1,54_ = 9.480, *p* < 0.005; Faith’s PD: *F*_1,22.4_ = 5.14, *p* = 0.033) (Figure 2; Table 1), whereas the Shannon diversity index did not differ between sexes.

**Figure 2:**
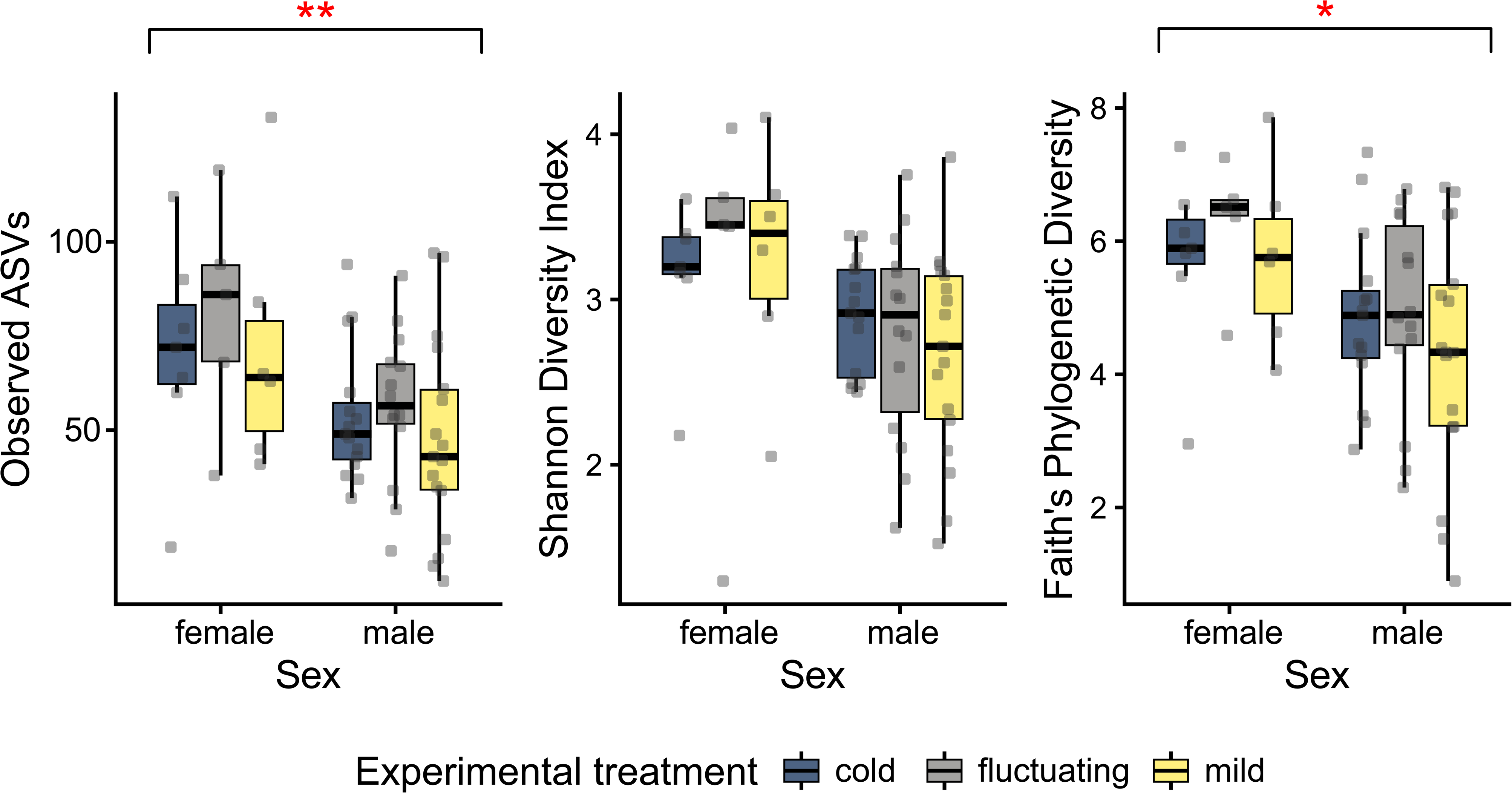
Gut microbiome diversity (Observed ASVs, Shannon diversity index, Faith’s Phylogenetic Diversity) among overwintering temperature treatment (cold, mild, fluctuating) and between sexes in common wall lizards. Boxplots show medians (centre line), interquartile ranges (boxes), and ranges (whiskers). Asterisks indicate significant differences among sexes (\**p* < 0.05, \*\**p* < 0.005).

**Table 1:**
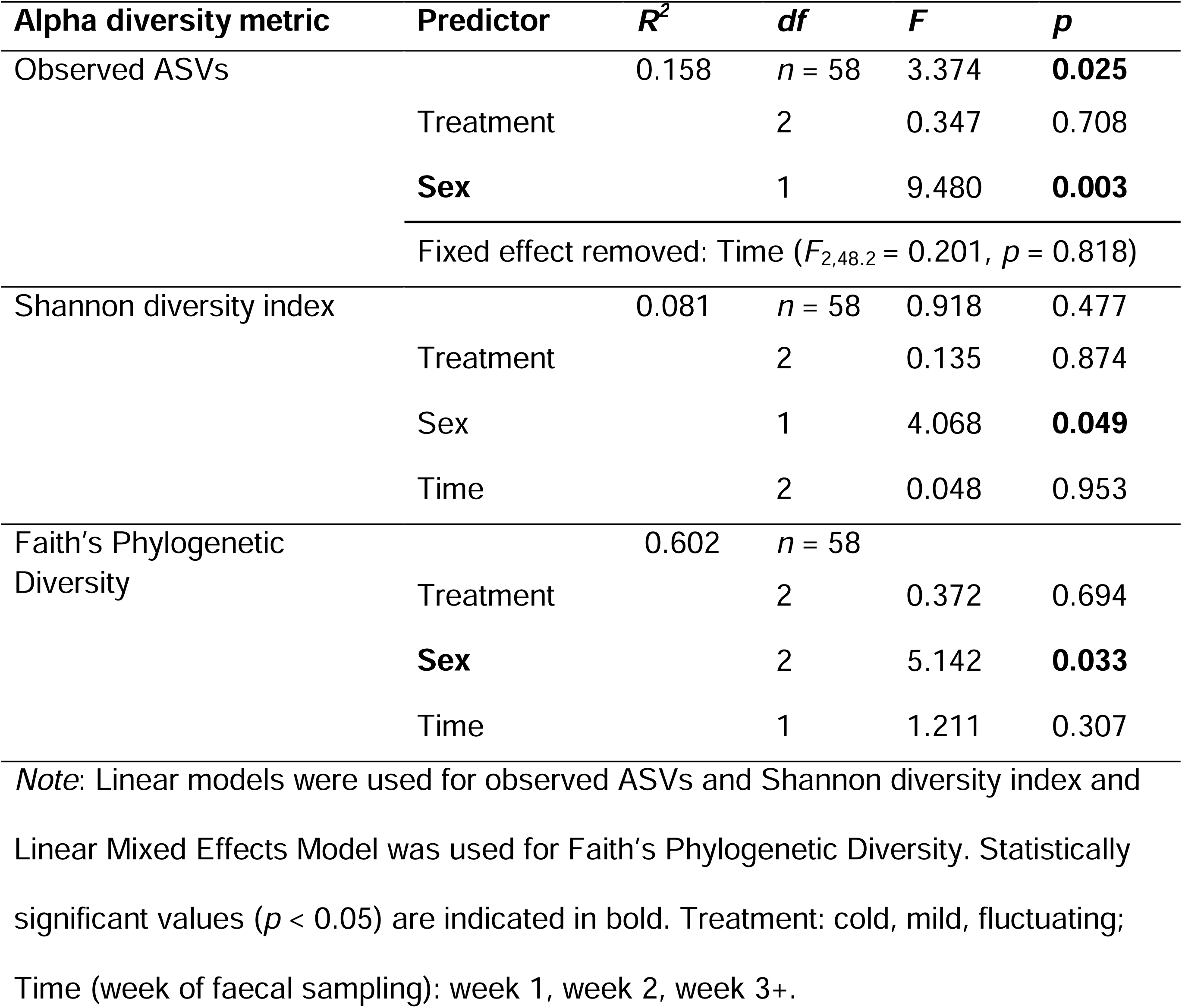
Association between gut microbiome diversity (Observed ASVs, Shannon diversity index, Faith’s Phylogenetic Diversity) and overwintering temperature treatments, sex, and sampling time points.

### 3.2 Gut microbiome composition varies across overwintering temperature, sex, and time

Overwintering temperature treatment was weakly associated with gut microbiome composition, explaining a small proportion of variance in both Bray-Curtis dissimilarities (*R*^2^ = 0.036) and unweighted UniFrac distances (*R*^2^ = 0.048; Table 2). Consistent with the small effect size, pairwise post-hoc comparisons among treatments were not significant (*p* > 0.05; Table S3) and PCoA ordinations showed substantial overlap among temperature groups (Figure 3a–b).

**Figure 3:**
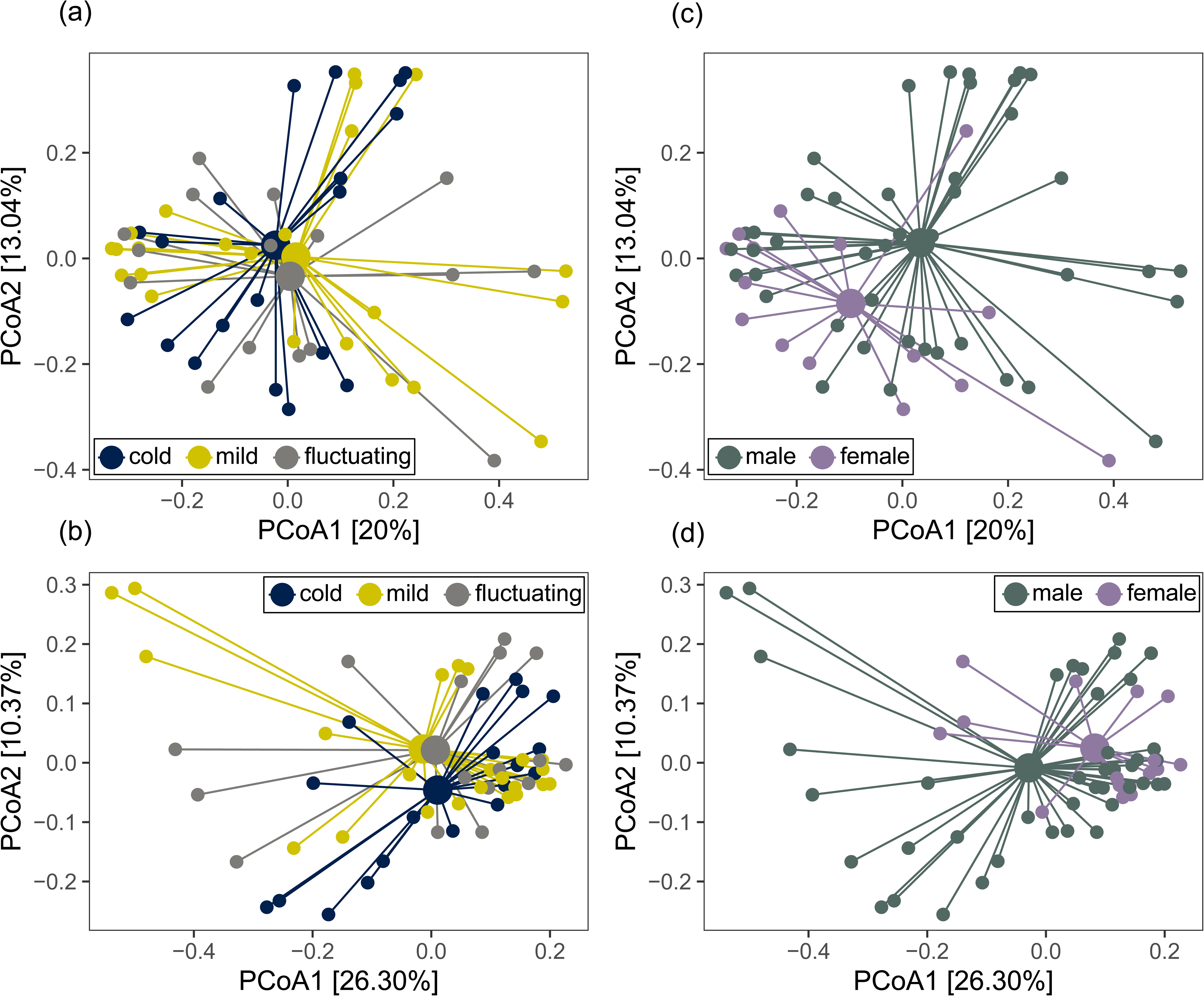
Shifts in gut microbiome composition of common wall lizards according to overwintering temperature treatment and sex. PCoA ordination was carried out using (a, c) Bray-Curtis distance matrices derived from ASV counts and (b, d) unweighted UniFrac distance matrices incorporating the phylogenetic relationships among ASVs. Each point represents a unique gut microbiome sample (*n* = 58 from 23 individuals). Large points indicate the group centroids, and lines connect samples to their respective group centroid. Percentages on axes indicate the proportion of variation explained by each principal coordinate.

**Table 2:**
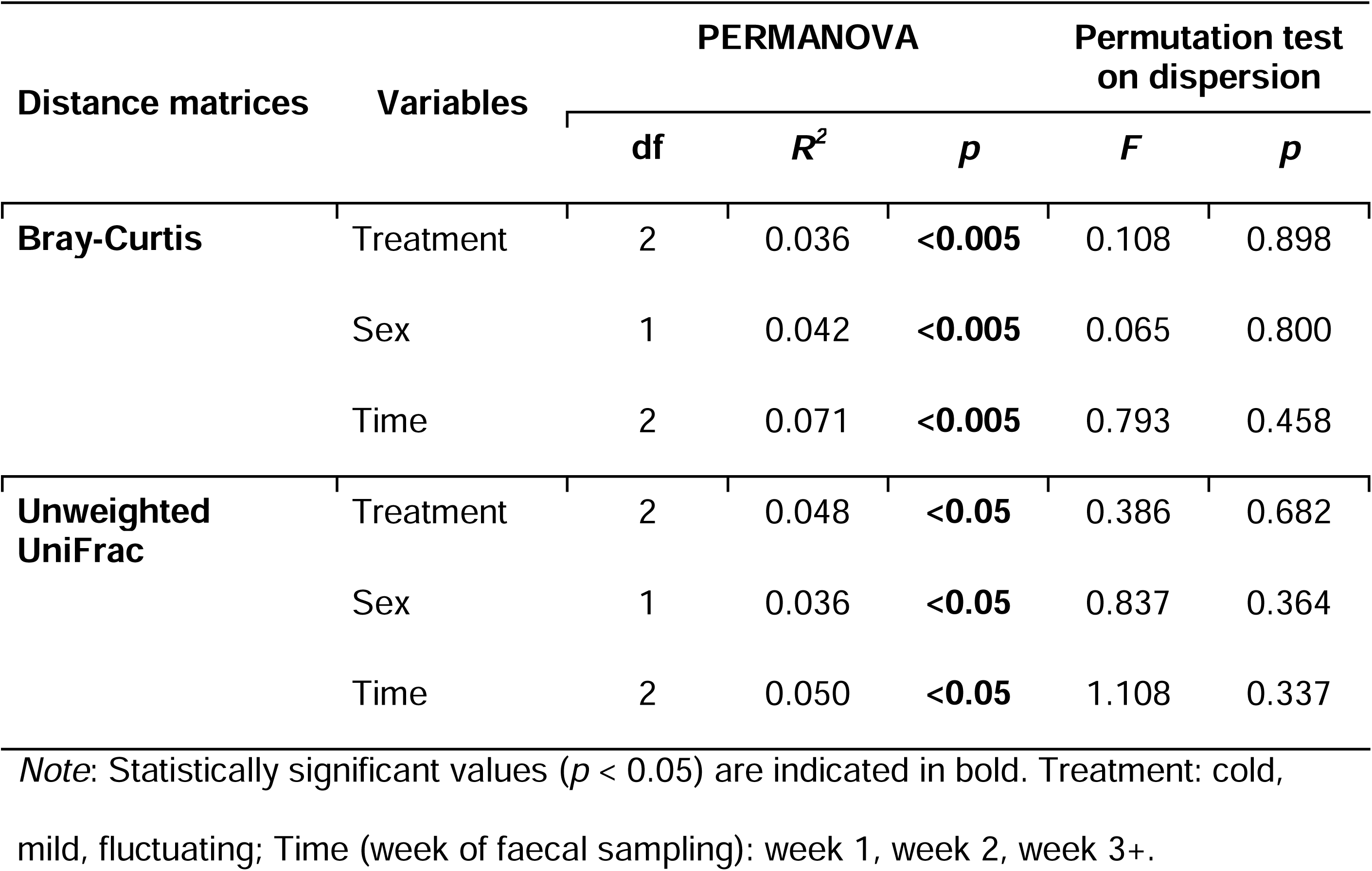
Results of permutational multivariate analysis of variance (PERMANOVA) tests and permutation tests of dispersion on beta diversity distances across overwintering temperature treatments, sex, and sampling time points. Permutations were restricted by individual ID.

Sex and sampling time explained comparable or greater variation in community composition than temperature treatment (Table 2). Microbiome composition differed between males and females (Bray-Curtis: *R*^2^ = 0.041, *p* < 0.005; unweighted UniFrac: *R*^2^ = 0.036, *p* < 0.05; Figure 3c–d) and varied across sampling time (Bray-Curtis: *R*^2^ = 0.071; unweighted UniFrac: *R*^2^ = 0.050; Figure S3). Samples from weeks 3+ showed greater separation in ordination space compared with earlier weeks (Figure S3; Table S3). Although significant, treatment, sex, and sampling time explained only ∼15% of the total variation in gut microbiome composition of common wall lizards.

Differential abundance analyses (DESeq2, adjusted *p* < 0.05) were performed at the ASV level but biological interpretation focused on the genus level. Only genera supported by at least two differentially abundant ASVs, or by a single ASV comprising ≥ 10% of the total community were retained. After filtering, 243 ASVs were kept and 262 low-support ASVs were excluded, yielding 23 genera belonging to five bacterial phyla (Actinomycetota, Bacillota, Bacteroidota, Fusobacteriota, and Pseudomonadota), including commonly detected members of the lizard gut microbiome (Figures 4 and S2).

**Figure 4:**
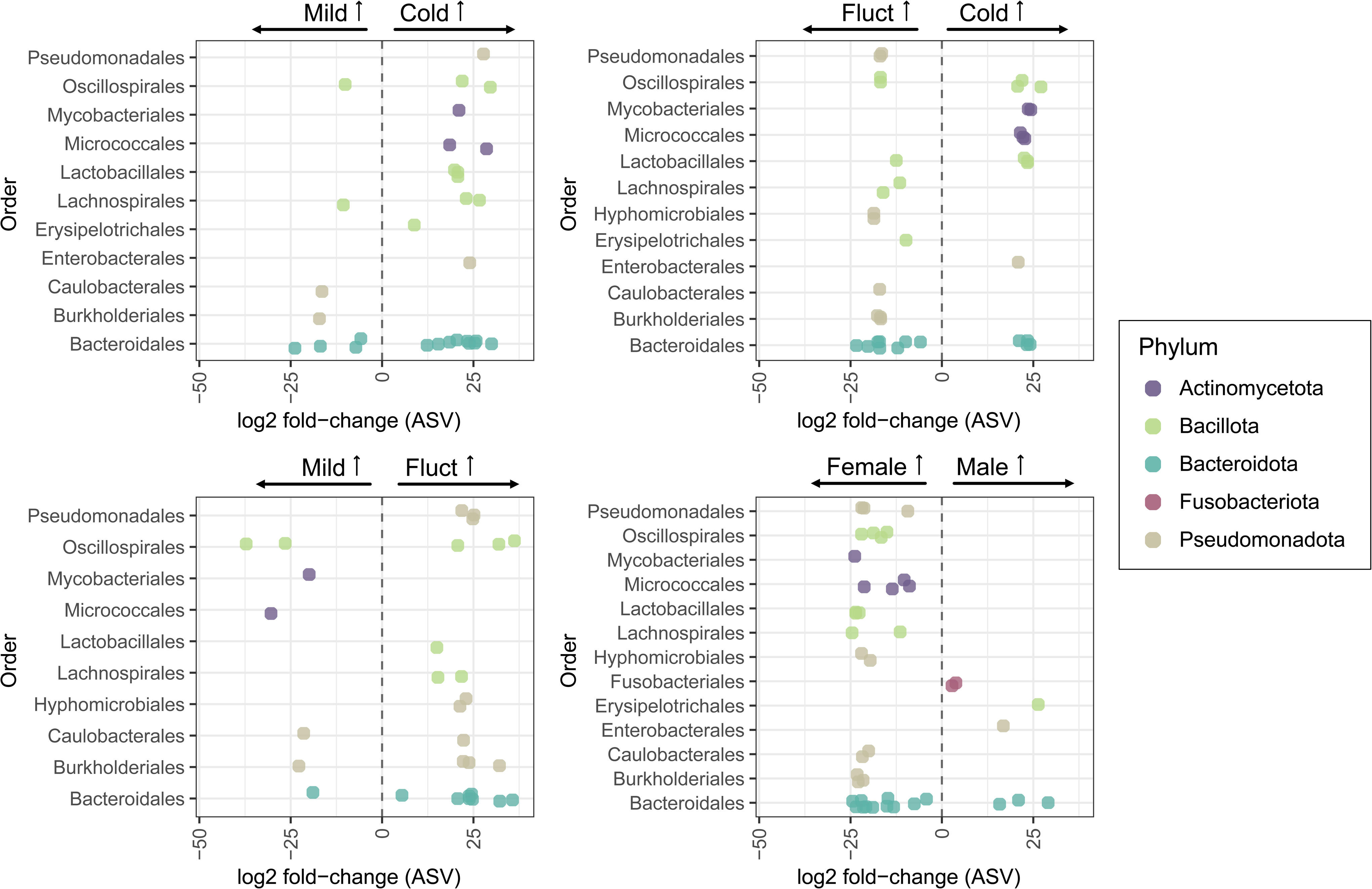
Differential abundance analysis (Deseq2) showing ASVs that differed significantly across temperature treatments (cold, mild, fluctuating) and sex (male vs female). Statistical testing was conducted at the ASV level using non-rarefied count data. For visualisation, significant ASVs are displayed at the order level to improve readability, with point colours indicating phylum affiliation. Each point represents log_2_ fold-change values of a significant ASV for a given contrast. Positive values indicate higher relative abundance in the right-hand group and negative values indicate higher relative abundance in the left-hand group. Multiple ASVs may belong to the same order; divergent log_2_ fold-change directions reflect heterogeneous responses among ASVs within a genus. Fluct = fluctuating temperature treatment.

Overwintering temperature treatment was associated with distinct shifts in the relative abundance of several genera, based on ASV-level differential abundance analysis (Figure S1). Compared to the cold and fluctuating treatments, the mild treatment enriched fewer taxa overall. In the cold and fluctuating treatments, most enriched taxa were also associated with fermentative metabolism (Koh et al., 2016; Louis and Flint, 2017), including short-chain fatty acid (SFCA) producers (e.g., *Roseburia*, *Intestinimonas*), protein and amino-acid fermenters (*Rikenella*, *Bacteroides*), and lactic acid bacteria (*Latilactobacillus*, *Pediococcus*). In contrast, the mild treatment enriched *Corynebacterium* and *Brevibacterium*, which include possibly opportunistic species (e.g., Mitchell and Markantonis, 2025; Panayiotou et al., 2025), compared to the fluctuating treatment, and also increased the abundance of *Odoribacter*, an SFCA producer, relative to the cold treatment.

Sex-specific differences in differential abundance were also detected (Figure S1). Females enriched a greater number of taxa, which were predominantly associated with fermentative and metabolic functions (e.g., *Roseburia, Pediococcus, Latilactobacillus, Bacteroides*), whereas genera enriched in males were largely putatively opportunistic or pathogenic, including *Hafnia-Obesumbacterium*, *Fusobacterium*, and *Butyricimonas* (Janda and Abbott, 2006; Li et al., 2025; Whitehill et al., 2024). Temporal contrasts further revealed restructuring of the gut microbiome, with later sampling weeks (weeks 3+) characterised by greater enrichment and dominance of fermenter genera compared to weeks 1 or 2, which showed greater enrichment of putatively opportunistic genera (e.g., *Hafnia-Obesumbacterium*, *Corynebacterium*, and *Brevibacterium*). Overall, overwintering temperatures, host sex, and time since emergence jointly shape gut microbiome composition, with cold and fluctuating treatments, female hosts, and later weeks following emergence showing greater enrichment of taxa linked to fermentative roles.

### 3.3 Overwintering temperature regimes induce microbiome dysbiosis

Community dispersion did not differ among overwintering temperature treatments for either Bray-Curtis (*F*_2,52_ = 0.16, *p* = 0.855) or unweighted UniFrac distances (*F*_2,52_ = 0.62, *p* = 0.542), indicating a comparable level of within-treatment heterogeneity (Figure 5a–b). However, a significant treatment×sex interaction was detected using Bray-Curtis dissimilarities (*F*_2,52_ = 3.81, *p* = 0.029) driven by greater dispersion among females than males in the cold treatment.

**Figure 5:**
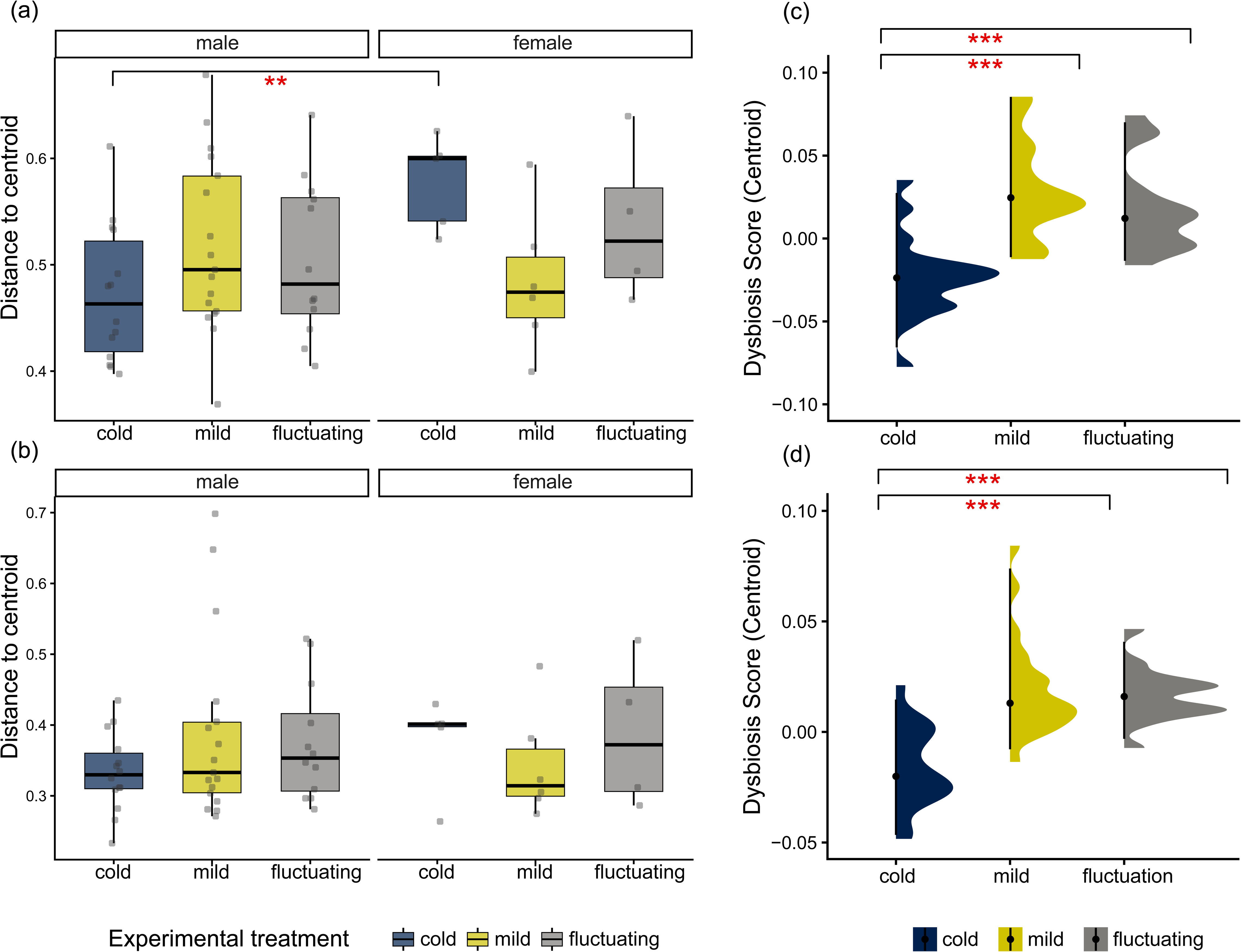
Effects of overwintering temperature and sex on dysbiosis metrics in common wall lizards. Left panels show differences in community dispersion among temperature treatments (cold, mild, fluctuating) and between sexes, calculated using *betadisperser* based on (a) Bray-Curtis distance matrices and (b) unweighted UniFrac distance matrices. Boxplots show medians (centre line), interquartile ranges (boxes), and ranges (whiskers). Right panels show dysbiosis scores across temperature treatments, defined as each sample’s Euclidean distance to the centroid of the cold control community, calculated using (c) Bray-Curtis distance matrices and (d) unweighted UniFrac distance matrices. Asterisks indicate significant differences among contrast groups (\*\**p* < 0.005, \*\*\**p* < 0.001).

We next quantified directional shifts in microbiome composition using a dysbiosis score defined as each sample’s Euclidean distance to the centroid of the cold control community. Dysbiosis scores differed significantly among overwintering temperature treatments for both Bray-Curtis (*F*_2,54_ = 13.97, *p* < 0.001) and unweighted UniFrac distances (*F*_2,54_ = 14.66, *p* < 0.001), with higher scores observed in the mild and fluctuating treatments relative to the cold treatment (Figure 5c–d; Table S4). For both community dispersion and directional deviation metrics, the area under the curve (AUC) values exceeded 0.8 for all pairwise contrasts, indicating strong discriminatory power of the dysbiosis metrics used (Figure S4).

## 4. Discussion

Climate warming is increasingly recognised as a major driver of change in animal-associated microbiomes (Bestion et al., 2017; Greenspan et al., 2020), but most empirical work has focused on microbiome dynamics during the active season. By contrast, the overwintering period, characterised by reduced metabolic activity, prolonged fasting, and altered host physiology, remains comparatively underexplored despite its potential importance for host-microbiome interactions. In overwintering vertebrates, gut microbiomes naturally exhibit seasonal restructuring, reflecting sensitivity to winter physiological states (Bunker et al., 2022; Liukkonen et al., 2024; Zhu et al., 2024). Winter warming may therefore represent an important but understudied source of host-microbiome disruption.

Here, we experimentally demonstrate that even modest alterations in overwintering temperature (i.e., 4°C difference) can reshape gut microbial communities in a temperate-zone ectotherm, the common wall lizard (*Podarcis muralis*). Rather than altering microbiome alpha diversity, winter warming was associated with subtle but consistent shifts in community composition and indicators of disruption in microbiome structure under warming conditions. These effects were further modulated by host sex and sampling time following emergence, highlighting the importance of intrinsic host traits and post-hibernation temporal dynamics in mediating microbiome responses to winter thermal environments. We discuss these findings by first addressing microbiome diversity responses, then compositional restructuring under different winter-warming regimes, and finally the role of sex and post-emergence recovery dynamics in shaping microbiome stability.

Contrary to our predictions, overwintering temperature treatments did not affect gut microbiome alpha diversity in common wall lizards. This contrasts with several studies reporting reduced microbial diversity in response to warming in ectotherms during the active season, including lizards (Bestion et al., 2017; Zhang et al., 2022b), fish (Zhou et al., 2022), and salamanders (Fontaine et al., 2018). However, responses of microbiome richness to warming are context-dependent, with increases reported in crocodile lizards (*Shinisaurus crocodilurus*; Lin et al., 2023) and corals (McDevitt-Irwin et al., 2019), and no detectable changes observed in tadpoles (Kohl and Yahn, 2016) and sea urchins (Schwob et al., 2025). Together, these contrasting patterns suggest that microbiome diversity responses to thermal change depend on host physiology, life-history stage, and likely the seasonal context in which warming occurs, as well as external factors such as the experimental design and conditions (Li et al., 2023).

A key distinction between our study and most previous work is that warming was applied during the overwintering period rather than during the host activity period. Overwintering ectotherms undergo profound physiological suppression, characterised by reduced metabolic rate, feeding activity, and gut motility, all of which likely slow microbial turnover and dampen competitive interactions within the gut (Olsson et al., 2025). Such conditions may buffer microbiome richness against moderate thermal perturbations even when relative abundances and community composition shift. Consistent with this, our thermal treatments were intentionally conservative and designed to reflect ecologically realistic winter conditions. More pronounced effects on alpha diversity may occur under the more extreme winter-warming scenarios projected for natural systems (Lee et al., 2023). Alternatively, stability in alpha diversity may reflect compensatory turnover, whereby losses of cold-adapted intestinal commensals under warmer conditions are offset by the proliferation of thermophile taxa. Together, these mechanisms possibly explain why overwintering microbiomes may be compositionally responsive, yet diversity-stable, under moderate warming.

In contrast to alpha diversity, overwintering temperatures shifted gut microbiome composition. Although effect sizes were modest, our two winter-warming regimes produced distinct compositional patterns. The constant mild temperature treatment was associated with limited enrichment of bacterial taxa, including *Corynebacterium* and *Brevibacterium*. Although these genera are well known for their fermentative role (e.g., in cheese production), several species have also been reported as sources of pathogenic infections in clinical isolates and in humans (Mitchell and Markantonis, 2025; Panayiotou et al., 2025). Their enrichment under constant mild warming may therefore reflect early or subtle shifts toward microbiome imbalance and potential for increased disease risk, rather than adaptive functional restructuring. However, further work is needed to confirm whether these changes translate into detrimental health outcomes.

In contrast, the fluctuating temperature regime, similar to the cold control treatment, was associated with enrichment of a broader range of bacterial taxa, relative to the mild treatment. These differentially abundant ASVs predominantly belonged to genera linked to fermentative metabolism (e.g., *Roseburia*, *Intestinimonas*, and *Rikenella*), playing an important role in energy extraction, short-chain fatty acid (SFCA) production, and gut homeostasis (Koh et al., 2016; Louis and Flint, 2017). This mirrors an enrichment in anaerobes during overwintering observed in some mammalian hibernators (Carey et al., 2013; Popov et al., 2025; Sommer et al., 2016), suggestive of an adaptive shift in response to torpor and fasting. This contrast between warming regimes suggests that thermal variability, rather than constant mild warming, may better preserve functionally beneficial microbial groups during overwintering. Because our fluctuating treatment included more cold days than warm days, the opposite thermal regime (i.e., more warm days than cold) could produce different outcomes and merits further investigation. Given projections that climate warming will increase not only mean winter temperatures but also thermal variability and the frequency of warm spells (Intergovernmental Panel on Climate Change, 2014; Lee et al., 2023), these findings additionally highlight the importance of examining multiple winter-warming patterns concurrently when assessing host-microbiome responses to climate change.

Beyond compositional shifts, winter warming was also associated with directional changes in overall microbiome structure. In both winter-warming regimes, we detected increased dysbiosis scores, quantified as directional deviation from the cold-treatment centroid using Euclidean distance (Shetty, 2022; Wei et al., 2021). Importantly, this increase was measured relative to the cold treatment, which itself represents an experimentally imposed reference rather than a true natural baseline. Consequently, our results should be interpreted as evidence that warmer overwintering conditions shift gut microbiome composition away from that observed under colder conditions, rather than as unequivocal evidence of pathological dysbiosis. Although the absolute magnitude of divergence was modest (∼0.05), the high discriminatory performance of the dysbiosis metric (AUC > 0.8) indicates that these shifts were consistent and detectable across individuals, suggesting biologically meaningful disruption of microbial communities (Huh and Roh, 2020; Jacobs et al., 2024). Despite these directional shifts, winter warming did not increase interindividual variability or overall community dispersion, providing no support for the Anna Karenina Principle (AKP), which predicts increased microbiome stochasticity and between-host variability under stress (Ma, 2020; Zaneveld et al., 2017). This finding indicates that winter warming, at least at the magnitude tested here, does not act as a strongly destabilising or chaotic stressor for the gut microbiome. Instead, microbiome responses appear structured and predictable, consistent with anti-AKP scenarios in which environmental stressors induce deterministic and stable community shifts. Such patterns have been reported in plants (Arnault et al., 2023), and in corals under heat stress (Zhu et al., 2023), as well as in systems shaped by strong stabilising interactions or environmental filters, such as polymicrobial infections with tightly integrated consortia (Kirst et al., 2015) or nutrientl1llimiting conditions such as iron deficiency (Pereira et al., 2015).

While winter warming did not increase stochasticity overall, patterns of microbiome dispersion differed between host sexes. The generality of dysbiosis as a universal microbial response to stress remains debated (Ma, 2020, 202; Olesen and Alm, 2016; Zaneveld et al., 2017), and certain stressors appear far more likely to trigger AKP-type increases in variability than others. For instance, psychological and social stressors acting through the gut-brain axis consistently induce AKP-like destabilisation (Garcia et al., 2025). In contrast, thermal stress during overwintering may operate through more constrained physiological pathways, such as suppressed metabolism, reduced microbial turnover, or limited nutrient flux, which could favour predictable, filtered community responses rather than stochastic divergence. We did, however, detect a sexl1lspecific pattern in community dispersion, with females in the cold group displaying higher microbiome community dispersion than males. Although this effect should be interpreted cautiously given samplel1lsize limitations, it suggests that sex may modulate microbiome stability under colder overwintering conditions. Such differences could arise from sexl1lspecific variation in metabolic rate, energy reserves, or immune function (e.g., Curlis et al., 2021; Hakim Saad and Shoukrey, 1988; Jönsson et al., 2009; Klein and Flanagan, 2016; Tobler et al., 2011).

Consistent with these dispersion patterns, sex emerged as a strong predictor of gut microbiome structure in common wall lizards. Females exhibited higher ASV richness and phylogenetic diversity than males, despite unequal sample sizes (n = 15 vs. 43), as well as a distinct community composition. Similar patterns have been observed from laboratory rodents, where males consistently exhibited lower alpha diversity (Tanelian et al., 2024), as well as in striped plateau lizards (Bunker et al., 2022). Such dimorphism can potentially be attributed to the bidirectional relationship between sex hormones and the gut microbiome (Jašarević et al., 2016), as well as sex-specific differences in physiological state and immune function. Additionally, sex shaped the pattern of differentially abundant taxa. Females showed enrichment of a greater number of genera, predominantly associated with fermentative metabolism (e.g., *Roseburia*, *Rickenella, Pediococcus, Lactilobacillus*). In contrast, males showed enrichment of fewer genera, which have been described in other hosts as opportunistic or potentially pathogenic (e.g., *Hafnia-Obesumbacterium*, *Fusobacterium*, and *Butyricimonas*; Janda and Abbott, 2006; Li et al., 2025; Wessendorf et al., 2024; Whitehill et al., 2024). Together, these findings suggest that sex-specific physiological responses to overwintering conditions may shape microbiome composition and stability in complex and sex-dependent ways.

Sampling time following emergence also appeared as an important factor shaping gut microbiome composition. Microbiomes sampled during weeks 1 and 2 were relatively similar and exhibited fewer differentially abundant taxa compared to week 3 onwards. Moreover, the enriched taxa predominantly belonged to genera described in other hosts as opportunistic or putatively pathogenic (e.g., *Hafnia-Obesumbacterium*, *Corynebacterium*, *Brevibacterium*, and *Brevibacterium*). From 3 weeks post-emergence and onwards, community composition diverged from earlier weeks and was characterised by a greater number of differentially abundant taxa, predominantly associated with fermentative metabolism (e.g., *Roseburiai, Rickenella, Alistipes,* and *Anaerotruncus*). This temporal pattern suggests that overwintering effects on the gut microbiome persist beyond emergence, consistent with a gradual post-hibernation recovery of host physiology. Overwintering induces immunosuppression (Bouma et al., 2010; Bouma et al., 2011), increasing susceptibility to opportunistic infections and parasitic burden (Ferguson et al., 2018b; Holden et al., 2021; Rumschlag and Boone, 2018), which may in turn favour non-beneficial gut taxa and delay recolonisation by commensal bacteria (Schlomann and Parthasarathy, 2019). In addition, temperature shifts can act directly on the microbial community, destabilising it and creating opportunities for colonisation by opportunistic taxa. Although later time points were pooled to increase statistical power, limiting temporal resolution, our results suggest that microbiome recovery following overwintering is gradual and extends several weeks beyond emergence (e.g., Lee et al., 2024). The delayed return to gut homeostasis probably reflects the combined effects of resumption of feeding, increased metabolic activity, and immune and physiological reactivation after prolonged fasting and torpor (Carey et al., 2013; Park and Do, 2024).

Due to limited sample size, we were unable to robustly test interactive effects such as investigating how treatment effects vary or dissipate over time, but future work could assess how recovery trajectories vary with overwintering conditions. Our findings are further constrained by the absence of pre-overwintering microbiome data, which prevents direct assessment of community shifts across the full overwintering period and post-emergence. Additionally, the lack of fitness or health-related outcomes limits our ability to evaluate the functional consequences of altered winter-warming patterns on host-microbiome dynamics. Nevertheless, our study provides rare experimental evidence that overwintering represents a biologically sensitive window during which climate warming can restructure host-associated microbiomes in ectotherms. By demonstrating that microbiome responses differ between constant and fluctuating warming regimes and are further modulated by sex and post-emergence timing, our findings suggest that the effects of winter conditions on gut microbiome dynamics are context-dependent rather than uniformly destabilising but have the potential for significant downstream consequences. Future work integrating functional, physiological, and fitness-related metrics will be essential to determine whether winter-induced microbiome shifts have lasting consequences for host performance and population resilience in a warming world.

## Data Accessibility Statement

Raw sequencing data, analysis datasets, metadata, and R scripts supporting this study are available in Zenodo at https://doi.org/10.5281/zenodo.18615422 under restricted access and will be made publicly available upon acceptance.

## Ethics approval statement

Animals were captured with permission from Bournemouth City Council and in partnership with the Amphibian and Reptile Conservation Trust. Experimental procedures and sample collection were carried out under Home Office Project Licence PP8703194, held by K.J.M.

## Author Contributions

M.R., K.J.M. and A.E. conceived the ideas and designed the methodology; M.R. collected the data; M.R., A.O. and A.E. analysed the data; M.R. and K.J.M. led the writing of the manuscript. All authors contributed critically to the drafts and gave final approval for publication.

## Acknowledgements

We thank the Bangor University animal technical team, including Rhys Morgan, Becca Snell, Andreas Akathiotis, Amy Battye and Mike Hayle for assistance in maintaining the lizard population. We also thank James Hicks and Benjamin Owens for helpful advice on lizard husbandry. Field efforts, including lizard capture, were greatly supported by Nick Moulton, from the Amphibian and Reptile Conservation Trust. We are also grateful to Anna Wood for valuable guidance during laboratory work, and to Owen Osborne for comments on molecular analysis.

## References

Almblad, H., Randall, T. E., Liu, F., Leblanc, K., Groves, R. A., Kittichotirat, W., Winsor, G. L., Fournier, N., Au, E., Groizeleau, J., et al. (2021). Bacterial cyclic diguanylate signaling networks sense temperature. Nat Commun 12, 1986.

Altizer, S., Ostfeld, R. S., Johnson, P. T. J., Kutz, S. and Harvell, C. D. (2013). Climate change and infectious diseases: from evidence to a predictive framework. Science 341, 514–519.

Amer, A., Spears, S., Vaughn, P. L., Colwell, C., Livingston, E. H., Mcqueen, W., Schill, A., Reichard, D. G., Gangloff, E. J. and Brock, K. M. (2024). Physiological phenotypes differ among color morphs in introduced common wall lizards (*Podarcis muralis*). Integrative Zoology 19, 505–523.

Arnault, G., Mony, C. and Vandenkoornhuyse, P. (2023). Plant microbiota dysbiosis and the Anna Karenina Principle. Trends in Plant Science 28, 18–30.

Barroso, F. M. (2023). Husbandry guidelines for the safe brumation of two lacertid lizard species (*Iberolacerta monticola* and *Podarcis lusitanicus*) in laboratory conditions v2. protocols.io.

Bates, D., Mächler, M., Bolker, B. and Walker, S. (2015). Fitting Linear Mixed-Effects Models Using lme4. Journal of Statistical Software 67, 1–48.

Benjamini, Y. and Hochberg, Y. (1995). Controlling the False Discovery Rate: A practical and powerful approach to multiple testing. Royal Statistical Society. Journal. Series B: Methodological 57, 289–300.

Bestion, E., Jacob, S., Zinger, L., Di Gesu, L., Richard, M., White, J. and Cote, J. (2017). Climate warming reduces gut microbiota diversity in a vertebrate ectotherm. Nat Ecol Evol 1, 0161.

Blix, A. S. (2016). Adaptations to polar life in mammals and birds. Journal of Experimental Biology 219, 1093–1105.

Bonnet, X., Bradshaw, D. and Shine, R. (1998). Capital versus income breeding: An ectothermic perspective. Oikos 83, 333–342.

Bouma, H. R., Carey, H. V. and Kroese, F. G. M. (2010). Hibernation: the immune system at rest? Journal of Leukocyte Biology 88, 619–624.

Bouma, H. R., Kroese, F. G. M., Kok, J. W., Talaei, F., Boerema, A. S., Herwig, A., Draghiciu, O., van Buiten, A., Epema, A. H., van Dam, A., et al. (2011). Low body temperature governs the decline of circulating lymphocytes during hibernation through sphingosine-1-phosphate. Proceedings of the National Academy of Sciences 108, 2052–2057.

Bray, J. R. and Curtis, J. T. (1957). An ordination of the upland forest communities of southern Wisconsin. Ecological Monographs 27, 326–349.

Brewer, S. M., Twittenhoff, C., Kortmann, J., Brubaker, S. W., Honeycutt, J., Massis, L. M., Pham, T. H. M., Narberhaus, F. and Monack, D. M. (2021). A *Salmonella* Typhi RNA thermosensor regulates virulence factors and innate immune evasion in response to host temperature. PLOS Pathogens 17, e1009345.

Bunker, M. E., Arnold, A. E. and Weiss, S. L. (2022). Wild microbiomes of striped plateau lizards vary with reproductive season, sex, and body size. Sci Rep 12, 20643.

Burke, R. L. and Ner, S. E. (2005). Seasonal and diel activity patterns of Italian wall lizards, *Podarcis sicula campestris*, in New York. Northeastern Naturalist 12, 349–360.

Callahan, B. J., McMurdie, P. J., Rosen, M. J., Han, A. W., Johnson, A. J. A. and Holmes, S. P. (2016). DADA2: High-resolution sample inference from Illumina amplicon data. Nat Methods 13, 581–583.

Carey, H. V., Walters, W. A. and Knight, R. (2013). Seasonal restructuring of the ground squirrel gut microbiota over the annual hibernation cycle. Am J Physiol Regul Integr Comp Physiol 304, R33–R42.

Chun, J. L., Ji, S. Y., Lee, S. D., Lee, Y. K., Kim, B. and Kim, K. H. (2020). Difference of gut microbiota composition based on the body condition scores in dogs. J Anim Sci Technol 62, 239–246.

Comizzoli, P., Power, M. L., Bornbusch, S. L. and Muletz-Wolz, C. R. (2021). Interactions between reproductive biology and microbiomes in wild animal species. Anim Microbiome 3, 87.

Curlis, J. D., Cox, C. L. and Cox, R. M. (2021). Sex-specific population differences in resting metabolism are associated with intraspecific variation in sexual size dimorphism of brown anoles. Physiological and Biochemical Zoology 94, 205–214.

Faith, D. P. (1992). Conservation evaluation and phylogenetic diversity. Biological Conservation 61, 1–10.

Ferguson, L. V., Dhakal, P., Lebenzon, J. E., Heinrichs, D. E., Bucking, C. and Sinclair, B. J. (2018a). Seasonal shifts in the insect gut microbiome are concurrent with changes in cold tolerance and immunity. Functional Ecology 32, 2357–2368.

Ferguson, L. V., Kortet, R. and Sinclair, B. J. (2018b). Eco-immunology in the cold: the role of immunity in shaping the overwintering survival of ectotherms. Journal of Experimental Biology 221, jeb163873.

Foa, A., Tosini, G. and Avery, R. (1992). Seasonal and diel cycles of activity in the ruin lizard, Podarcis sicula. Herpetological Journal 2, 86–89.

Fontaine, S. S., Novarro, A. J. and Kohl, K. D. (2018). Environmental temperature alters the digestive performance and gut microbiota of a terrestrial amphibian. J Exp Biol 221, jeb187559.

Garcia, I., Kilic, F., Bryan, C.-A., Castro-Vildosola, J., Jonnalagadda, S. A., Kasturi, A., Tilly, J., Smith, J., Valentin, S., Moncayo, S., et al. (2025). Social stress changes gut microbiome composition in male, female, and aggressor mice. Brain, Behavior, & Immunity - Health 50, 101138.

Greenspan, S. E., Migliorini, G. H., Lyra, M. L., Pontes, M. R., Carvalho, T., Ribeiro, L. P., Moura-Campos, D., Haddad, C. F. B., Toledo, L. F., Romero, G. Q., et al. (2020). Warming drives ecological community changes linked to host-associated microbiome dysbiosis. Nat. Clim. Chang. 10, 1057–1061.

Hakim Saad, A. and Shoukrey, N. (1988). Sexual dimorphism on the immune responses of the snake, *Psammophis sibilans*. Immunobiology 177, 404–419.

Hanning, I. and Diaz-Sanchez, S. (2015). The functionality of the gastrointestinal microbiome in non-human animals. Microbiome 3, 51.

Hartig, F. (2024). DHARMa: Residual Diagnostics for Hierarchical (Multi-Level / Mixed) Regression Models.

Hector, T. E., Hoang, K. L., Li, J. and King, K. C. (2022). Symbiosis and host responses to heating. Trends in Ecology & Evolution 37, 611–624.

Hoffbeck, C., Middleton, D. M. R. L., Nelson, N. J. and Taylor, M. W. (2023). 16S rRNA gene-based meta-analysis of the reptile gut microbiota reveals environmental effects, host influences and a limited core microbiota. Molecular Ecology 32, 6044–6058.

Hoffbeck, C., Middleton, D. M. R. L., Lamar, S. K., Keall, S. N., Nelson, N. J. and Taylor, M. W. (2024). Gut microbiome of the sole surviving member of reptile order Rhynchocephalia reveals biogeographic variation, influence of host body condition and a substantial core microbiota in tuatara across New Zealand. Ecology and Evolution 14, e11073.

Holden, K. G., Gangloff, E. J., Gomez-Mancillas, E., Hagerty, K. and Bronikowski, A. M. (2021). Surviving winter: Physiological regulation of energy balance in a temperate ectotherm entering and exiting brumation. General and Comparative Endocrinology 307, 113758.

Huh, J.-W. and Roh, T.-Y. (2020). Opportunistic detection of *Fusobacterium nucleatum* as a marker for the early gut microbial dysbiosis. BMC Microbiol 20, 208.

Hylander, B. L. and Repasky, E. A. (2019). Temperature as a modulator of the gut microbiome: What are the implications and opportunities for thermal medicine? Int J Hyperthermia 36, 83–89.

Intergovernmental Panel on Climate Change. (2014). Climate Change 2013—the physical science basis: Working Group I Contribution to the Fifth Assessment Report of the Intergovernmental Panel on Climate Change. Cambridge: Cambridge University Press.

Jacobs, J. P., Spencer, E. A., Helmus, D. S., Yang, J. C., Lagishetty, V., Bongers, G., Britton, G., Gettler, K., Reyes-Mercedes, P., Hu, J., et al. (2024). Age-related patterns of microbial dysbiosis in multiplex inflammatory bowel disease families. Gut 73, 1953–1964.

Janda, J. M. and Abbott, S. L. (2006). The genus *Hafnia*: from soup to nuts. Clin Microbiol Rev 19, 12–28.

Jandhyala, S. M., Talukdar, R., Subramanyam, C., Vuyyuru, H., Sasikala, M. and Reddy, D. N. (2015). Role of the normal gut microbiota. World J Gastroenterol 21, 8787–8803.

Jašarević, E., Morrison, K. E. and Bale, T. L. (2016). Sex differences in the gut microbiome–brain axis across the lifespan. Philos Trans R Soc Lond B Biol Sci 371, 20150122.

Jönsson, K. I., Herczeg, G., O’Hara, R. B., Söderman, F., Ter Schure, A. F. H., Larsson, P. and Merilä, J. (2009). Sexual patterns of prebreeding energy reserves in the common frog *Rana temporaria* along a latitudinal gradient. Ecography 32, 831–839.

Kembel, S. W., Cowan, P. D., Helmus, M. R., Cornwell, W. K., Morlon, H., Ackerly, D. D., Blomberg, S. P. and Webb, C. O. (2010). Picante: R tools for integrating phylogenies and ecology. Bioinformatics 26, 1463–1464.

Kim, C., Alrefaei, R., Bushlaibi, M., Ndegwa, E., Kaseloo, P. and Wynn, C. (2019). Influence of growth temperature on thermal tolerance of leading foodborne pathogens. Food Sci Nutr 7, 4027–4036.

Kirst, M. E., Li, E. C., Alfant, B., Chi, Y.-Y., Walker, C., Magnusson, I. and Wang, G. P. (2015). Dysbiosis and alterations in predicted functions of the subgingival microbiome in chronic periodontitis. Appl Environ Microbiol 81, 783–793.

Klein, S. L. and Flanagan, K. L. (2016). Sex differences in immune responses. Nat Rev Immunol 16, 626–638.

Koh, A., De Vadder, F., Kovatcheva-Datchary, P. and Bäckhed, F. (2016). From dietary fiber to host physiology: Short-Chain Fatty Acids as key bacterial metabolites. Cell 165, 1332–1345.

Kohl, K. D. and Yahn, J. (2016). Effects of environmental temperature on the gut microbial communities of tadpoles. Environmental Microbiology 18, 1561–1565.

Kohl, K. D., Brun, A., Magallanes, M., Brinkerhoff, J., Laspiur, A., Acosta, J. C., Caviedes-Vidal, E. and Bordenstein, S. R. (2017). Gut microbial ecology of lizards: insights into diversity in the wild, effects of captivity, variation across gut regions and transmission. Mol Ecol 26, 1175–1189.

Kolenda, K., Skawiński, T., Majtyka, T., Majtyka, M., Kuśmierek, N., Natalia Kuśmierek, Starzecka, A. and Jablonski, D. (2020). Biology and origin of isolated north-easternmost populations of the common wall lizard, *Podarcis muralis*. Amphibia-Reptilia 41, 429–443.

Lee, H., Calvin, K., Dasgupta, D., Krinner, G., Mukherji, A., Thorne, P., Trisos, C., Romero, J., Aldunce, P., Barrett, K., et al. (2023). Climate change 2023: synthesis report. Contribution of working groups I, II and III to the sixth assessment report of the intergovernmental panel on climate change. The Australian National University.

Lee, J.-E., Park, J.-K. and Do, Y. (2024). Gut microbiome diversity and function during hibernation and spring emergence in an aquatic frog. PLOS ONE 19, e0298245.

Lesser, M. P., Fiore, C., Slattery, M. and Zaneveld, J. (2016). Climate change stressors destabilize the microbiome of the Caribbean barrel sponge, *Xestospongia muta*. Journal of Experimental Marine Biology and Ecology 475, 11–18.

Li, J., Bates, K. A., Hoang, K. L., Hector, T. E., Knowles, S. C. L. and King, K. C. (2023). Experimental temperatures shape host microbiome diversity and composition. Global Change Biology 29, 41–56.

Li, Z., Liu, J., Li, J., Zhou, Z., Huang, X., Gopinath, D., Luo, P., Wang, Q. and Shan, D. (2025). *Fusobacterium* in the microbiome: from health to disease across the oral–gut axis and beyond. npj Biofilms Microbiomes 11, 200.

Lin, Z., He, M., Zhong, C., Li, Y., Tang, S., Kang, X. and Wu, Z. (2023). Responses of gut microbiota in crocodile lizards (*Shinisaurus crocodilurus*) to changes in temperature. Front. Microbiol. 14, 1263917.

Liukkonen, M., Muriel, J., Martínez-Padilla, J., Nord, A., Pakanen, V.-M., Rosivall, B., Tilgar, V., van Oers, K., Grond, K. and Ruuskanen, S. (2024). Seasonal and environmental factors contribute to the variation in the gut microbiome: A large-scale study of a small bird. Journal of Animal Ecology 93, 1475–1492.

Louis, P. and Flint, H. J. (2017). Formation of propionate and butyrate by the human colonic microbiota. Environmental Microbiology 19, 29–41.

Love, M. I., Huber, W. and Anders, S. (2014). Moderated estimation of fold change and dispersion for RNA-seq data with DESeq2. Genome Biol 15, 550.

Lozupone, C. and Knight, R. (2005). UniFrac: a new phylogenetic method for comparing microbial communities. Appl Environ Microbiol 71, 8228–8235.

Ma, Z. (Sam). (2020). Testing the Anna Karenina Principle in human microbiome-associated diseases. iScience 23, 101007.

MacLeod, K. J., Kohl, K. D., Trevelline, B. K. and Langkilde, T. (2022). Context-dependent effects of glucocorticoids on the lizard gut microbiome. Molecular Ecology 31, 185–196.

Martin, M. (2011). Cutadapt removes adapter sequences from high-throughput sequencing reads. EMBnet.journal 17, 10–12.

Martin, M. O., Gilman, F. R. and Weiss, S. L. (2010). Sex-specific asymmetry within the cloacal microbiota of the striped plateau lizard, *Sceloporus virgatus*. Symbiosis 51, 97–105.

McDevitt-Irwin, J. M., Garren, M., McMinds, R., Vega Thurber, R. and Baum, J. K. (2019). Variable interaction outcomes of local disturbance and El Niño-induced heat stress on coral microbiome alpha and beta diversity. Coral Reefs 38, 331–345.

McFall-Ngai, M., Hadfield, M. G., Bosch, T. C. G., Carey, H. V., Domazet-Lošo, T., Douglas, A. E., Dubilier, N., Eberl, G., Fukami, T., Gilbert, S. F., et al. (2013). Animals in a bacterial world, a new imperative for the life sciences. Proceedings of the National Academy of Sciences 110, 3229–3236.

Michaelides, S., While, G. M., Bell, C. and Uller, T. (2013). Human introductions create opportunities for intra-specific hybridization in an alien lizard. Biological Invasions 15, 1101–1112.

Michaelides, S., While, G. M., Zajac, N. and Uller, T. (2015). Widespread primary, but geographically restricted secondary, human introductions of wall lizards, *Podarcis muralis*. Molecular Ecology 24, 2702–2714.

Mitchell, B. I. and Markantonis, J. E. (2025). An underestimated pathogen: *Corynebacterium* species. Journal of Clinical Microbiology 63, e01552–24.

Moeller, A. H., Ivey, K., Cornwall, M. B., Herr, K., Rede, J., Taylor, E. N. and Gunderson, A. R. (2020). The lizard gut microbiome changes with temperature and is associated with heat tolerance. Applied and Environmental Microbiology 86, e01181–20.

Moss, J. B. and MacLeod, K. J. (2022). A quantitative synthesis of and predictive framework for studying winter warming effects in reptiles. Oecologia 200, 259–271.

Oksanen, J., Simpson, G. L., Blanchet, F. G., Kindt, R., Legendre, P., Minchin, P. R., O’Hara, R. B., Solymos, P., Stevens, M. H. H., Szoecs, E., et al. (2025). vegan: Community Ecology Package.

Olesen, S. W. and Alm, E. J. (2016). Dysbiosis is not an answer. Nat Microbiol 1, 16228.

Olsson, C., Gräns, A. and Brijs, J. (2025). Beyond ecology: the importance of gut motility in predicting the responses of species to climate change. J Exp Biol 228, jeb249822.

Oskyrko, O., Laakkonen, H., Silva-Rocha, I., Jablonski, D., Marushchak, O., Uller, T. and Carretero, M. A. (2020). The possible origin of the common wall lizard, *Podarcis muralis* (Laurenti, 1768) in Ukraine. Herpetozoa 33, 87–93.

Panayiotou, T., Vasilopoulou, A., Baliou, S., Tsantes, A. G. and Ioannou, P. (2025). *Brevibacterium* species infections in humans—A narrative review. Microorganisms 13, 1097.

Paradis, E. and Schliep, K. (2019). ape 5.0: an environment for modern phylogenetics and evolutionary analyses in R. Bioinformatics 35, 526–528.

Park, J.-K. and Do, Y. (2024). Combined effect of seasons and life history in an anuran strengthens the response and relationship between their physiology and gut microbiota. Sci Rep 14, 10137.

Pereira, D. I. A., Aslam, M. F., Frazer, D. M., Schmidt, A., Walton, G. E., McCartney, A. L., Gibson, G. R., Anderson, G. J. and Powell, J. J. (2015). Dietary iron depletion at weaning imprints low microbiome diversity and this is not recovered with oral nano Fe(III). Microbiologyopen 4, 12–27.

Perry, C., Sarraude, T., Billet, M., Minot, E., Gangloff, E. J. and Aubret, F. (2024). Sex-dependent shifts in body size and condition along replicated elevational gradients in a montane colonising ectotherm, the common wall lizard (*Podarcis muralis*). Oecologia 206, 335–346.

Popov, I. V., Peshkova, D. A., Lukbanova, E. A., Tsurkova, I. S., Emelyantsev, S. A., Krikunova, A. A., Malinovkin, A. V., Chikindas, M. L., Ermakov, A. M. and Popov, I. V. (2025). Gut microbiota dynamics in hibernating and active *Nyctalus noctula*: Hibernation-associated loss of diversity and anaerobe enrichment. Veterinary Sciences 12, 559.

Quast, C., Pruesse, E., Yilmaz, P., Gerken, J., Schweer, T., Yarza, P., Peplies, J. and Glöckner, F. O. (2013). The SILVA ribosomal RNA gene database project: improved data processing and web-based tools. Nucleic Acids Res 41, D590–D596.

Raselimanana, M., Wüster, W., Blount, J. D., Mitchell, C. A., Morgan, R., Wilkinson, J. W. and MacLeod, K. J. (2025). Experimental winter warming increases activity with signs of potential DNA damage in common wall lizards. J Exp Biol 228, jeb251440.

Robin, X., Turck, N., Hainard, A., Tiberti, N., Lisacek, F., Sanchez, J.-C. and Müller, M. (2011). pROC: an open-source package for R and S+ to analyze and compare ROC curves. BMC Bioinformatics 12, 77.

Rocca, J. D., Simonin, M., Blaszczak, J. R., Ernakovich, J. G., Gibbons, S. M., Midani, F. S. and Washburne, A. D. (2019). The Microbiome Stress Project: Toward a global meta-analysis of environmental stressors and their effects on microbial communities. Front. Microbiol. 9, 3272.

Roncarati, D., Vannini, A. and Scarlato, V. (2025). Temperature sensing and virulence regulation in pathogenic bacteria. Trends in Microbiology 33, 66–79.

Ruiz-Rodríguez, M., Soler, J. J., Lucas, F. S., Heeb, P., José Palacios, M., Martín-Gálvez, D., De Neve, L., Pérez-Contreras, T., Martínez, J. G. and Soler, M. (2009). Bacterial diversity at the cloaca relates to an immune response in magpie *Pica pica* and to body condition of great spotted cuckoo *Clamator glandarius* nestlings. Journal of Avian Biology 40, 42–48.

Rumschlag, S. and Boone, M. (2018). High juvenile mortality in amphibians during overwintering related to fungal pathogen exposure. Dis. Aquat. Org. 131, 13–28.

Sacchi, R., Pupin, F., Gentilli, A., Rubolini, D., Scali, S., Fasola, M. and Galeotti, P. (2009). Male-male combats in a polymorphic lizard: residency and size, but not color, affect fighting rules and contest outcome. Aggress Behav 35, 274–283.

Sakelarieva, L., Pulev, A. and Mitrevichin, E. (2023). Winter activity of *Podarcis muralis* (Laurenti, 1768) and *P. erhardii* (Bedriaga, 1876) in urban and suburban habitats in the city of Blagoevgrad, Bulgaria. Preprints.

Salter, S. J., Cox, M. J., Turek, E. M., Calus, S. T., Cookson, W. O., Moffatt, M. F., Turner, P., Parkhill, J., Loman, N. J. and Walker, A. W. (2014). Reagent and laboratory contamination can critically impact sequence-based microbiome analyses. BMC Biol 12, 87.

Schlomann, B. H. and Parthasarathy, R. (2019). Timescales of gut microbiome dynamics. Current Opinion in Microbiology 50, 56–63.

Schloss, P. D. (2024). Rarefaction is currently the best approach to control for uneven sequencing effort in amplicon sequence analyses. mSphere 9, e00354-23.

Schulte, U., Gassert, F., Geniez, P., Veith, M. and Hochkirch, A. (2012). Origin and genetic diversity of an introduced wall lizard population and its cryptic congener. Amphibia-reptilia 33, 129–140.

Schulte-Hostedde, A., Zinner, B., Millar, J. and Hickling, G. (2005). Restitution of mass-size residuals: Validating body condition indices. Ecology 86, 155–163.

Schwob, G., Delleuze, M., Motreuil, S., Marschal, C., Saucède, T., Orlando, J., Poulin, E. and Cabrol, L. (2025). Gut microbiome plasticity and host resistance in response to ocean warming in sub-Antarctic sea urchins. BMC Biol 23, 343.

Sepulveda, J. and Moeller, A. H. (2020). The effects of temperature on animal gut microbiomes. Front. Microbiol. 11, 384.

Shannon, C. E. (1948). A mathematical theory of communication. The Bell System Technical Journal 27, 379–423.

Shetty, S. (2022). dysbiosisR: an R package for calculating microbiome dysbiosis measures.

Shi, N., He, M., Lin, Z., Kang, X. and Wu, Z. (2025). Climate warming induces age-specific gut microbiota alterations and functional metagenomic changes in captive crocodile lizards (*Shinisaurus crocodilurus*). Global Ecology and Conservation 63, e03905.

Shine, R. (2005). Life-history evolution in reptiles. Annual Review of Ecology, Evolution, and Systematics 36, 23–46.

Slevin, M. C., Houtz, J. L., Vitousek, M. N., Baldassarre, D. T. and Anderson, R. C. (2025). Ornamentation and body condition, but not glucocorticoids, predict wild songbird cloacal microbiome community and diversity. Oikos 2025, e10905.

Sommer, F., Ståhlman, M., Ilkayeva, O., Arnemo, J. M., Kindberg, J., Josefsson, J., Newgard, C. B., Fröbert, O. and Bäckhed, F. (2016). The gut microbiota modulates energy metabolism in the hibernating brown bear *Ursus arctos*. Cell Rep 14, 1655–1661.

Stothart, M. R., Bobbie, C. B., Schulte-Hostedde, A. I., Boonstra, R., Palme, R., Mykytczuk, N. C. S. and Newman, A. E. M. (2016). Stress and the microbiome: linking glucocorticoids to bacterial community dynamics in wild red squirrels. Biol Lett 12, 20150875.

Tanelian, A., Nankova, B., Miari, M. and Sabban, E. L. (2024). Microbial composition, functionality, and stress resilience or susceptibility: unraveling sex-specific patterns. Biol Sex Differ 15, 20.

Teyssier, A., Lens, L., Matthysen, E. and White, J. (2018). Dynamics of gut microbiota diversity during the early development of an avian host: Evidence from a cross-foster experiment. Front Microbiol 9, 1524.

Tobler, M., Healey, M., Wilson, M. and Olsson, M. (2011). Basal superoxide as a sex-specific immune constraint. Biol Lett 7, 906–908.

Tremaroli, V. and Bäckhed, F. (2012). Functional interactions between the gut microbiota and host metabolism. Nature 489, 242–249.

Turner, R. K. and Maclean, I. M. D. (2022). Microclimate-driven trends in spring-emergence phenology in a temperate reptile (*Vipera berus*): Evidence for a potential “climate trap”? Ecology and Evolution 12, e8623.

Vose, R. S., Russell S. Vose, Easterling, D. R. and Gleason, B. (2005). Maximum and minimum temperature trends for the globe: An update through 2004. Geophysical Research Letters 32, L23822.

Wei, S., Bahl, M. I., Baunwall, S. M. D., Hvas, C. L. and Licht, T. R. (2021). Determining gut microbial dysbiosis: a review of applied indexes for assessment of intestinal microbiota imbalances. Appl Environ Microbiol 87, e00395–21.

Wessendorf, L., Brezina, T., Dusse, F., Wiegel, P. and Boschert, A. L. (2024). *Butyricimonas faecihominis*: an atypically resistant bacterium implicated in abscess formation - a case report. BMC Infect Dis 24, 697.

While, G. M., Williamson, J., Joseph Williamson, Prescott, G. W., Horváthová, T., Fresnillo, B., Fresnillo, B., Belén Fresnillo, Beeton, N. J., Halliwell, B., et al. (2015). Adaptive responses to cool climate promotes persistence of a non-native lizard. Proceedings of The Royal Society B: Biological Sciences 282, 20142638–20142638.

Whitehill, G., Zhuo, R. and Yang, S. (2024). *Butyricimonas paravirosa* bacteremia associated with acute terminal ileitis: Case report and literature review. Anaerobe 90, 102918.

Williams, C. M., Hérault, B., Henry, H. A. L. and Sinclair, B. J. (2015). Cold truths: how winter drives responses of terrestrial organisms to climate change. Biological Reviews 90, 214–235.

Wilsterman, K., Ballinger, M. A. and Williams, C. M. (2021). A unifying, eco-physiological framework for animal dormancy. Functional Ecology 35, 11–31.

Zaneveld, J. R., Mcminds, R. and Vega Thurber, R. (2017). Stress and stability: applying the Anna Karenina principle to animal microbiomes. Nature Microbiology 2, 17121.

Zhang, Z., Tang, H., Chen, P., Xie, H. and Tao, Y. (2019). Demystifying the manipulation of host immunity, metabolism, and extraintestinal tumors by the gut microbiome. Sig Transduct Target Ther 4, 41.

Zhang, L., Yang, F., Li, T., Dayananda, B., Lin, L. and Lin, C. (2022a). Lessons from the diet: Captivity and sex shape the gut microbiota in an oviparous lizard (*Calotes versicolor*). Ecology and Evolution 12, e8586.

Zhang, Z., Zhu, Q., Chen, J., Khattak, R. H., Li, Z., Teng, L. and Liu, Z. (2022b). Insights into the composition of gut microbiota in response to environmental temperature: The case of the Mongolia racerunner (*Eremias argus*). Global Ecology and Conservation 36, e02125.

Zhou, J., Nelson, T. M., Rodriguez Lopez, C., Sarma, R. R., Zhou, S. J. and Rollins, L. A. (2020). A comparison of nonlethal sampling methods for amphibian gut microbiome analyses. Mol Ecol Resour 20, 844–855.

Zhou, C., Yang, S., Ka, W., Gao, P., Li, Y., Long, R. and Wang, J. (2022). Association of gut microbiota with metabolism in rainbow trout under acute heat stress. Front. Microbiol. 13, 846336.

Zhu, W., Wang, H., Li, X., Liu, X., Zhu, M., Wang, A. and Li, X. (2023). Consistent responses of coral microbiome to acute and chronic heat stress exposures. Marine Environmental Research 185, 105900.

Zhu, X., Jiang, N., Mai, T., Wu, S., Yao, Y., Du, Y., Lin, C., Lin, L. and Ji, X. (2024). Gut microbial communities are seasonally variable in warm-climate lizards hibernating in the winter months. Microorganisms 12, 1974.

