## Supplementary Materials for "Winter warming shapes gut microbiome composition and directional dysbiosis in a temperate lizard"

MOLECULAR ECOLOGY

MOLECULAR ECOLOGY

### SUPPLEMENTARY METHODS

#### 1. DNA extraction, 16S rRNA amplification and sequencing

Prior to processing the faecal samples collected, we conducted a validation test to compare the efficiency of two extraction kits commonly used for microbiome analyses: QIAamp Fast DNA stool mini kit (QIAGEN, Venlo, The Netherlands) and DNeasy Powersoil Pro kit (QIAGEN, Venlo, The Netherlands). For the validation test, we used four lizard faecal samples collected opportunistically that we processed immediately after collection. Each sample was split into two equal subsamples after manually removing the uric acid fraction (the white portion) with sterile forceps. Uric acids are known to inhibit enzymatic reactions such as PCR (Khan et al., 1991). Faecal subsamples were extracted with both kits following the manufacturer's instructions. DNA concentration and purity (A260/280) were quantified using a NanoDrop Lite Plus spectrophotometer (ThermoFisher Scientific, Waltham, MA, USA). The DNeasy PowerSoil Pro Kit consistently produced higher DNA concentrations and better purity ratios, so we used that kit for all subsequent extractions, including the control substrate samples. DNA was eluted in 50 µL of elution buffer to maximise DNA concentration. All DNA extracts were stored at −20 °C until PCR amplification and library preparation. To monitor potential contamination introduced during the extraction process, one extraction blank (containing no biological material) was included in each batch and processed identically to the samples.

The V4 region of the 16S rRNA gene (~253 bp) was amplified using primers 515F_Parada (5′-GTGYCAGCMGCCGCGGTAA-3′; Parada et al., 2016) and 806R_Apprill (5′-GGACTACNVGGGTWTCTAAT-3′; Apprill et al., 2015) in a one-step PCR. 16S rRNA gene amplicon data are widely used for describing reptile gut microbiota (Hoffbeck et al., 2023). To enable sample identification, forty-eight 12 bp multiplex identifier (MID) tags were incorporated into the forward and reverse primers. Unique dual indexes were used for each sample to ensure that any erroneously tagged reads, such as from PCR chimeras or tag-jumping during library preparation (Rodriguez-Martinez et al., 2023), would be removed during de-multiplexing.

PCR reactions (25 µL) contained 12.5 µL Q5 Hot Start High-Fidelity 2X Master Mix (NEB, Ipswich, MA, USA), 1 µL of each primer (10µM), 2 µL of template DNA, and 8.5 µL of ultrapure water. Cycling conditions were: (1) an initial denaturation at 98 °C for 1 min; followed by (2) 35 cycles of 98 °C for 15 s, 50 °C for 30 s, and 72 °C for 30 s; and (3) a final extension at 72 °C for 10 min. Each PCR batch included a blank and a PCR negative control to monitor potential contamination, and substrate control samples were also amplified alongside faecal samples. PCR products were quantified using Qubit Broad Range DNA assays (ThermoFisher Scientific, Waltham, MA, USA) and visualised by electrophoresis on 1.5% TAE agarose gels. Libraries were pooled equimolarly based on Qubit-quantified concentrations, purified with Agencourt AMPure XP magnetic beads (Fisher Scientific, Waltham, MA, USA), and sequenced with Illumina NovaSeq 2 x 250 bp at Novogene (Beijing, China).

### 2. Bioinformatics

Raw paired-end Illumina reads were concatenated and demultiplexed by sample-specific MID tags using *cutadapt* v5.1 (Martin, 2011). Quality assessment and sequence processing were conducted using the DADA2 pipeline v1.8 (Callahan et al., 2016) following default recommended parameters for quality filtering, denoising, merging of paired-end reads, and chimera removal using the consensus method. Sequences falling outside the expected amplicon length for the V4 region (200-300 bp) were discarded.

Taxonomic assignment of ASVs was performed using DADA2's naïve Bayesian classifier (Callahan et al., 2016) trained on the SILVA 16S reference database (v138.2; Quast et al., 2013). We removed low abundance ASVs (≤ 10 total reads), ASVs lacking phylum-level classification, and non-bacterial sequences (chloroplasts, mitochondria). A phylogenetic tree was generated from aligned ASV sequences using *DECIPHER* v3.2.0 (Wright, 2016), optimized by maximum-likelihood in *phangorn* v2.12.1 (Schliep, 2011), and midpoint-rooted for phylogenetic metrics. We combined the resulting ASV table with metadata, taxonomy table, and phylogenetic tree using the *phyloseq* package v1.50.0 (McMurdie and Holmes, 2013). Potential contaminants were identified with the *decontam* package v1.26.0 (prevalence method, threshold = 0.1; Davis et al., 2018) based on extraction blanks, PCR negatives, and substrate controls, resulting in the removal of 13 ASVs.

Following filtering, sequencing depth per sample ranged from 5,521 to 102,334 reads (mean = 37,196; median = 34,550). Rarefaction curves were generated using *vegan* v2.7 (Oksanen et al., 2025) and indicated that ASV richness approached an asymptote around 8,000 reads for the majority of samples (see rarefaction curves in Supplementary Figure S5). We therefore rarefied the ASV tables to an even depth of 8,000 reads prior to diversity analyses following the recommendations of Schloss (2024). This rarefaction depth balanced retention of samples with robust estimation of diversity metrics. Rarefaction was applied for diversity-based analysis (alpha and beta diversity metrics); all other analyses were conducted on non-rarefied data. After quality control and removal of nonbacterial sequences, sequencing yielded a total of 2,975,669 reads (mean ± SE: 37,196 ± 1979 reads per sample), grouped into a total of 615 ASVs across 80 samples, with rarefaction retaining 503 ASVs across 78 samples.

### 3. References

**Apprill, A., McNally, S., Parsons, R. and Weber, L.** (2015). Minor revision to V4 region SSU rRNA 806R gene primer greatly increases detection of SAR11 bacterioplankton. *Aquat. Microb. Ecol.* **75**, 129–137.

**Callahan, B. J., McMurdie, P. J., Rosen, M. J., Han, A. W., Johnson, A. J. A. and Holmes, S. P.** (2016). DADA2: High-resolution sample inference from Illumina amplicon data. *Nat Methods* **13**, 581–583.

**Davis, N. M., Proctor, D. M., Holmes, S. P., Relman, D. A. and Callahan, B. J.** (2018). Simple statistical identification and removal of contaminant sequences in marker-gene and metagenomics data. *Microbiome* **6**, 226.

**Hoffbeck, C., Middleton, D. M. R. L., Nelson, N. J. and Taylor, M. W.** (2023). 16S rRNA gene-based meta-analysis of the reptile gut microbiota reveals environmental effects, host influences and a limited core microbiota. *Molecular Ecology* **32**, 6044–6058.

**Khan, G., Kangro, H. O., Coates, P. J. and Heath, R. B.** (1991). Inhibitory effects of urine on the polymerase chain reaction for cytomegalovirus DNA. *J Clin Pathol* **44**, 360–365.

**Martin, M.** (2011). Cutadapt removes adapter sequences from high-throughput sequencing reads. *EMBnet.journal* **17**, 10–12.

**McMurdie, P. and Holmes, S. P.** (2013). phyloseq: An R package for reproducible interactive analysis and graphics of microbiome census data. *PLOS One* **8**, e61217.

**Oksanen, J., Simpson, G. L., Blanchet, F. G., Kindt, R., Legendre, P., Minchin, P. R., O'Hara, R. B., Solymos, P., Stevens, M. H. H., Szoecs, E., et al.** (2025). vegan: Community Ecology Package.

**Parada, A. E., Needham, D. M. and Fuhrman, J. A.** (2016). Every base matters: assessing small subunit rRNA primers for marine microbiomes with mock communities, time series and global field samples. *Environmental Microbiology* **18**, 1403–1414.

**Quast, C., Pruesse, E., Yilmaz, P., Gerken, J., Schweer, T., Yarza, P., Peplies, J. and Glöckner, F. O.** (2013). The SILVA ribosomal RNA gene database project: improved data processing and web-based tools. *Nucleic Acids Res* **41**, D590–D596.

**Rodriguez-Martinez, S., Klaminder, J., Morlock, M. A., Dalén, L. and Huang, D. Y.-T.** (2023). The topological nature of tag jumping in environmental DNA metabarcoding studies. *Molecular Ecology Resources* **23**, 621–631.

**Schliep, K. P.** (2011). phangorn: phylogenetic analysis in R. *Bioinformatics* **27**, 592–593.

MOLECULAR ECOLOGY

**Schloss, P. D.** (2024). Rarefaction is currently the best approach to control for uneven sequencing effort in amplicon sequence analyses. *mSphere* **9**, e00354-23.

**Wright, E., S.** (2016). Using DECIPHER v2.0 to Analyze Big Biological Sequence Data in R. *The R Journal* **8**, 352–359.

### SUPPLEMENTARY RESULTS

#### 1. Overwintering temperature does not affect interindividual variability

We quantified interindividual variability as the mean pairwise distance between samples within each group using Bray-Curtis and unweighted UniFrac distances. Interindividual variability did not differ among overwintering temperature treatments for either Bray-Curtis ($X_2^2$ = 0.52, *p* = 0.772) or unweighted UniFrac distances ($X_2^2$ = 1.60, *p* = 0.450; Table S4).

#### 2. Host body condition is not correlated with microbiome diversity or dysbiosis

Spearman's rank correlations revealed no significant associations between body condition index at emergence (BCI) and any alpha diversity metrics (ASV richness, Shannon diversity index, Faith's phylogenetic diversity) or dysbiosis metrics, including community dispersion, directional deviation from the cold control community, and interindividual variability (Table S5). This pattern was consistent in both females (*n* = 15) and males (*n* = 43), with all FDR-corrected *p* ≥ 0.05.

MOLECULAR ECOLOGY

### SUPPLEMENTARY TABLES

**Table S1:** Summary of faecal sampling effort across treatments (cold, fluctuating, mild winter temperature), sexes, and weekly time points (week 1, week 2, week 3+)

| Treatment | Sex | | Time points | | | Total |
|---|---|---|---|---|---|---|
| | M | F | Week 1 | Week 2 | Week 3+ | |
| Cold | 15 | 7 | 7 | 8 | 8 | 26 |
| Fluctuating | 14 | 5 | 7 | 11 | 6 | 27 |
| Mild | 19 | 6 | 6 | 13 | 7 | 27 |
| | | | | | | 80 |

**Table S2:** Post hoc pairwise comparisons testing differences in gut microbiome alpha diversity (Observed ASVs, Shannon diversity index, and Faith's Phylogenetic Diversity) across overwintering temperature treatments, host sex, and post-emergence sampling time points.

| Alpha diversity metric | Predictor | Contrast | Estimates | *t* | *p* |
|---|---|---|---|---|---|
| **Observed ASVs** | Treatment | cold-mild | 2.12 | 0.28 | 0.958 |
| | | cold-fluctuating | -4.48 | -0.54 | 0.852 |
| | | mild-fluctuating | -6.61 | -0.83 | 0.687 |
| **Shannon diversity index** | Treatment | cold-mild | 0.10 | 0.50 | 0.807 |
| | | cold-fluctuating | 0.08 | 0.37 | 0.929 |
| | | mild-fluctuating | -0.02 | -0.10 | 0.994 |
| | Time | week1-week2 | 0.02 | 0.01 | 0.993 |
| | | week1-week3+ | 0.07 | 0.31 | 0.950 |
| | | week2-week3+ | 0.04 | 0.21 | 0.975 |

MOLECULAR ECOLOGY

| | | | | | |
|---|---|---|---|---|---|
| **Faith's Phylogenetic Diversity** | Treatment | cold-mild | 0.28 | 0.43 | 0.905 |
| | | cold-fluctuating | -0.26 | -0.39 | 0.921 |
| | | mild-fluctuating | -0.54 | -0.86 | 0.673 |
| | Time | week1-week2 | 0.07 | 0.19 | 0.980 |
| | | week1-week3+ | -0.50 | -1.17 | 0.478 |
| | | week2-week3+ | -0.560 | -1.494 | 0.304 |

*Note*: Treatment: cold, mild, fluctuating; Time (week of faecal sampling): week 1, week 2, week 3+.

**Table S3:** Full PERMANOVA results including fixed effects, interaction terms, and pairwise comparisons, for beta diversity distances across overwintering temperature treatments, sex, and sampling time points. Permutations were restricted by individual ID.

| **Distance matrices** | **Predictor** | **PERMANOVA** | | |
|---|---|---|---|---|
| | | **df** | $R^2$ | ***p*** |
| **Bray-Curtis** | **Treatment** | 2 | 0.036 | **<0.005** |
| | **Sex** | 1 | 0.042 | **<0.005** |
| | **Time** | 2 | 0.071 | **<0.005** |
| | Treatment x sex | 2 | 0.051 | 0.134 |
| | Treatment x time | 4 | 0.063 | 0.518 |
| **Unweighted UniFrac** | **Treatment** | 2 | 0.048 | **<0.05** |
| | **Sex** | 1 | 0.036 | **<0.05** |
| | **Time** | 2 | 0.050 | **<0.05** |
| | Treatment x sex | 2 | 0.037 | 0.193 |
| | Treatment x time | 4 | 0.076 | 0.106 |

MOLECULAR ECOLOGY

| Distance matrices | Predictor | Pairs | Pairwise PERMANOVA | |
|---|---|---|---|---|
| | | | $R^2$ | $p$ |
| **Bray-Curtis** | Treatment | cold-mild | 0.030 | 0.217 |
| | | cold-fluctuating | 0.025 | 0.601 |
| | | mild-fluctuating | 0.024 | 0.485 |
| | Time | week1-week2 | 0.032 | 0.201 |
| | | week1-week3+ | 0.086 | **0.001** |
| | | week2-week3+ | 0.052 | **0.015** |
| **Unweighted UniFrac** | Treatment | cold-mild | 0.038 | 0.106 |
| | | cold-fluctuating | 0.042 | 0.137 |
| | | mild-fluctuating | 0.031 | 0.275 |
| | Time | week1-week2 | 0.013 | 0.934 |
| | | week1-week3+ | 0.051 | 0.059 |
| | | week2-week3+ | 0.051 | **0.040** |

*Note*: LizardID was included as a blocking factor to control for repeated sampling of individuals. Statistically significant values ($p < 0.05$) are indicated in bold. Treatment: cold, mild, fluctuating; Time (week of faecal sampling): week 1, week 2, week 3+.

**Table S4:** Dysbiosis metrics (community dispersion, directional shift, and interindividual variability) with post-hoc comparisons

| BRAY-CURTIS DISTANCE | | | | |
|---|---|---|---|---|
| **Dysbiosis metric** | **Variables** | **df** | $F/\mathcal{X}^2$ | $p$ |
| Community dispersion (betadisperser) | Treatment | 2 | 0.16 | 0.855 |
| | Sex | 1 | 1.99 | 0.164 |
| | **Treatment x sex** | 2 | 3.81 | **0.029** |

MOLECULAR ECOLOGY

| | | | | |
|---|---|---|---|---|
| Directional deviation from the control cold treatment (Euclidean distance) | **Treatment** | 2 | 20.65 | **<0.001** |
| | Sex | 1 | 0.60 | 0.441 |
| | Interaction removed: Treatment x sex ($F_{2,52}$ = 1.04, $p$ = 0.359) | | | |
| Interindividual variability | Treatment | 2 | 0.52 | 0.772 |

**UNWEIGHTED UNIFRAC DISTANCE**

| **Dysbiosis metric** | **Variables** | **df** | ***F/$\mathcal{X}^2$*** | ***p*** |
|---|---|---|---|---|
| Community dispersion | Treatment | 2 | 0.62 | 0.542 |
| | Sex | 1 | 0.03 | 0.868 |
| | Treatment x sex | 2 | 0.91 | 0.409 |
| Directional deviation | **Treatment** | 2 | 21.98 | **<0.001** |
| | Sex | 1 | 0.03 | 0.866 |
| | Interaction removed: Treatment x sex ($F_{2,52}$ = 1.67, $p$ = 0.199) | | | |
| Interindividual variability | Treatment | 2 | 1.60 | 0.450 |

**BRAY-CURTIS DISTANCE (Post-hoc comparisons)**

| **Dysbiosis metrics** | **Predictor** | | **Contrast** | **Estimate** | ***t*** | ***p*** |
|---|---|---|---|---|---|---|
| Community dispersion | Sex | Female | cold-mild | 0.09 | 2.17 | 0.086 |
| | | | cold-fluctuating | 0.04 | 0.84 | 0.680 |
| | | | mild-fluctuating | -0.05 | -1.16 | 0.482 |
| | | Male | cold-mild | -0.05 | -1.75 | 0.196 |
| | | | cold-fluctuating | -0.03 | -1.17 | 0.479 |
| | | | mild-fluctuating | 0.01 | 0.46 | 0.889 |

| | | | | | | |
|---|---|---|---|---|---|---|
| Community dispersion | Treatment | Cold | male-female | -0.11 | -2.8 | **0.007** |
| | | Mild | male-female | 0.03 | 0.99 | 0.326 |
| | | Fluctuating | male-female | -0.03 | -0.78 | 0.437 |
| Directional deviation | Treatment | | cold-mild | -0.05 | -6.85 | **<0.001** |
| | | | cold-fluctuating | -0.04 | -5.48 | **<0.001** |
| | | | mild-fluctuating | -0.01 | 0.81 | 0.700 |
| Interindividual variability | Treatment | | cold-mild | -0.001 | -0.02 | 1.000 |
| | | | cold-fluctuating | -0.02 | -0.62 | 1.000 |
| | | | mild-fluctuating | -0.01 | -0.63 | 1.000 |

**UNWEIGHTED UNIFRAC DISTANCE (Post-hoc comparisons)**

| Dysbiosis metrics | Predictor | | Contrast | Estimate | *t* | *p* |
|---|---|---|---|---|---|---|
| Community dispersion | Sex | Female | cold-mild | 0.10 | 0.71 | 0.757 |
| | | | cold-fluctuating | -0.01 | -0.05 | 0.998 |
| | | | mild-fluctuating | -0.11 | -0.72 | 0.751 |
| | | Male | cold-mild | -0.12 | -1.42 | 0.336 |
| | | | cold-fluctuating | -0.11 | -1.19 | 0.462 |
| | | | mild-fluctuating | 0.01 | 0.12 | 0.992 |
| | Treatment | Cold | male-female | -0.13 | -1.05 | 0.300 |
| | | Mild | male-female | 0.09 | 0.85 | 0.402 |
| | | Fluctuating | male-female | -0.03 | -0.19 | 0.849 |
| Directional deviation | Treatment | | cold-mild | -0.04 | -6.14 | **<0.001** |
| | | | cold-fluctuating | -0.03 | -5.24 | **<0.001** |
| | | | mild-fluctuating | 0.002 | 0.38 | 0.922 |

| | | | | | |
|---|---|---|---|---|---|
| Interindividual variability | Treatment | cold-mild | -0.04 | -1.09 | 0.873 |
| | | cold-fluctuating | -0.04 | -1.14 | 0.810 |
| | | mild-fluctuating | -0.003 | -0.10 | 1.000 |

*Note:* Statistically significant values ($p < 0.05$) are indicated in bold. Treatment: cold, mild, fluctuating

**Table S5:** Correlation coefficients and p-values for relationships between body condition index, gut microbiome alpha diversity (ASV richness, Shannon diversity index, Faith's phylogenetic diversity), and dysbiosis metrics (community dispersion, directional deviation from the cold control community, and interindividual variability)

| | | Body condition index at emergence | | | |
|---|---|---|---|---|---|
| | | Females | | Males | |
| **Alpha diversity metrics** | | ***r*** | ***p*** | ***r*** | ***p*** |
| Observed ASV | | 0.29 | 0.529 | 0.17 | 0.665 |
| Shannon diversity | | 0.10 | 0.711 | 0.001 | 0.993 |
| Faith's phylogenetic diversity | | 0.36 | 0.529 | 0.16 | 0.665 |
| **Dysbiosis metrics** | | ***r*** | ***p*** | ***r*** | ***p*** |
| Community dispersion | Bray-Curtis | -0.29 | 0.529 | 0.03 | 0.993 |
| | Unweighted UniFrac | -0.33 | 0.529 | 0.04 | 0.993 |
| Directional deviation | Bray-Curtis | -0.16 | 0.710 | -0.33 | 0.229 |
| | Unweighted UniFrac | -0.16 | 0.710 | 0.30 | 0.229 |
| Interindividual variability | Bray-Curtis | -0.23 | 0.629 | 0.07 | 0.993 |
| | Unweighted UniFrac | -0.32 | 0.529 | -0.004 | 0.993 |

*Note*: Spearman's rank correlations were performed and p-values were corrected using the Benjamini-Hochberg FDR method

MOLECULAR ECOLOGY

### SUPPLEMENTARY FIGURES

**Fig. S1:** Differential abundance analysis (Deseq2) showing ASVs that differed significantly across temperature treatments (cold, mild, fluctuating), sex (male vs female), and sampling time (week of faecal sampling). Each point represents a single ASV, plotted at the genus the genus level for visualisation, with point colours indicating phylum affiliation. Positive $\log_2$ fold-change values indicate higher relative abundance in the right-hand group of contrast and negative values indicate higher relative abundance in the left-hand group. Multiple ASVs from the same genus may show divergent responses, reflecting heterogeneity within genera. Fluct = fluctuating temperature treatment.

**Fig. S2:** Relative abundance of bacterial phyla in the gut microbiome of common wall lizards. (a) shows mean relative abundances across overwintering temperature treatments (cold, mild, fluctuating) and (b) shows mean relative abundances between sexes.

**Fig. S3:** Shifts in gut microbiome composition of common wall lizards according to sampling time (week of faecal sampling). PCoA ordination was carried out using (a) Bray-Curtis dissimilarities from ASV counts and (b) unweighted UniFrac distances incorporating the phylogenetic relationships among ASVs. Each point represents a unique gut microbiome sample ($n$ = 58 from 23 individuals). Large points indicate the group centroids, and lines connect samples to their respective group centroid. Percentages on axes indicate the proportion of variation explained by each principal coordinate.

**Fig. S4:** Area Under the Receiver Operating Characteristic Curve (AUC-ROC) evaluating the discriminatory performance of dysbiosis metrics. (a) AUC-ROC based on Bray-Curtis distances comparing community dispersion between females and males in the cold temperature treatment; (b, d) AUC-ROC based on Bray-Curtis distances assessing directional deviation in the mild and fluctuating winter temperature treatments relative to the control cold; (c, e) AUC-ROC based on unweighted UniFrac distances assessing directional deviation in the mild and fluctuating winter temperature treatments relative to the control cold. All AUC values exceed 0.8, indicating strong discriminatory power of the dysbiosis metrics.

**Fig. S5:** Rarefaction curves of gut microbiome diversity in common wall lizards (*Podarcis muralis*) across overwintering temperature treatments. Each line represents a sample, showing observed ASV richness as a function of sequencing depth. Lines are coloured by treatments. The dashed vertical line at 8,000 reads indicates the sequencing depth used for rarefaction prior to downstream analyses, based on asymptote trends. Curves indicate that sequencing depth captured most microbial diversity in the majority of the samples.
