## Supplementary figures and images for "Winter warming shapes gut microbiome composition and directional dysbiosis in a temperate lizard"

### Supplementary Figure S1

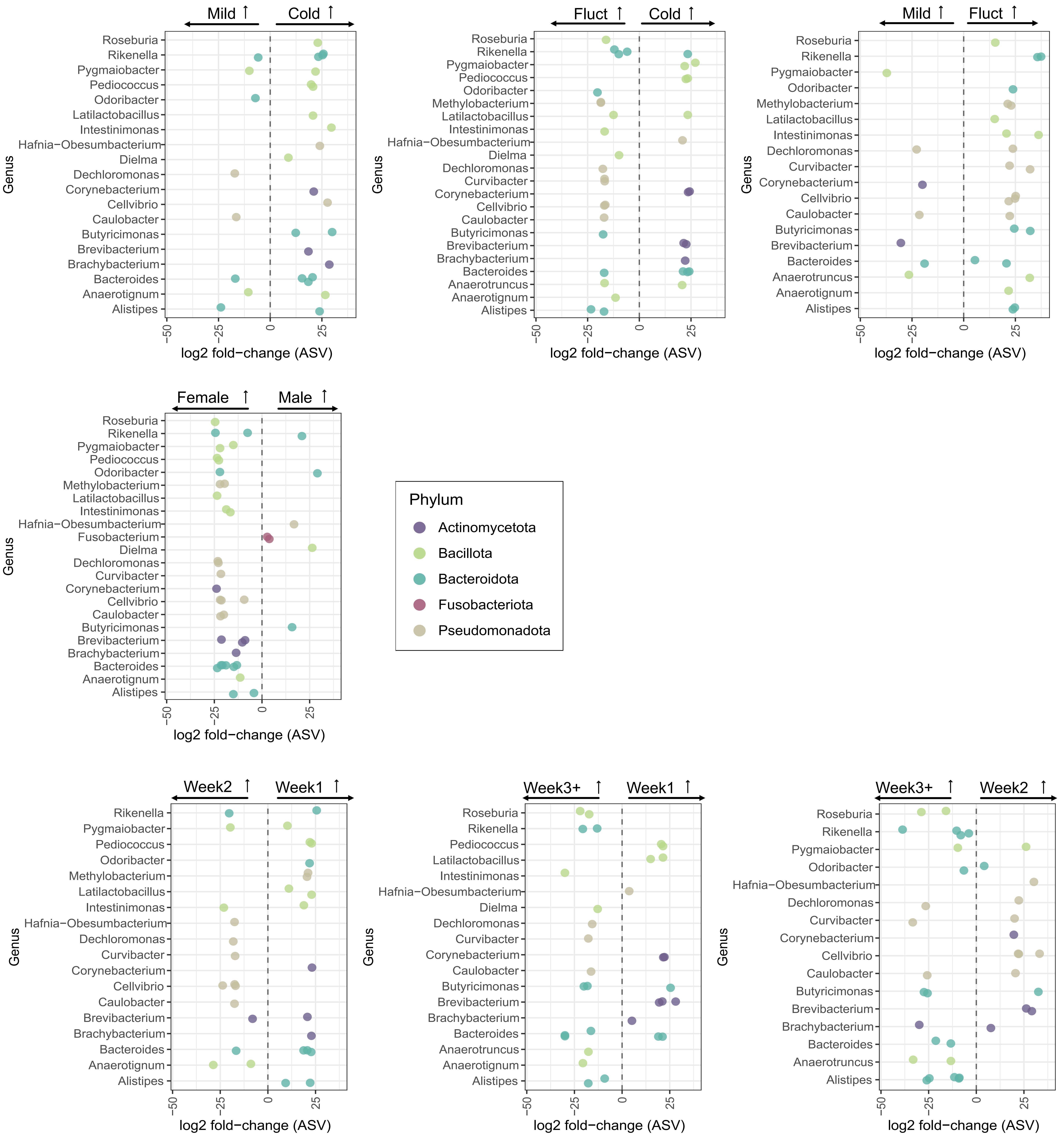

### Supplementary Figure S2

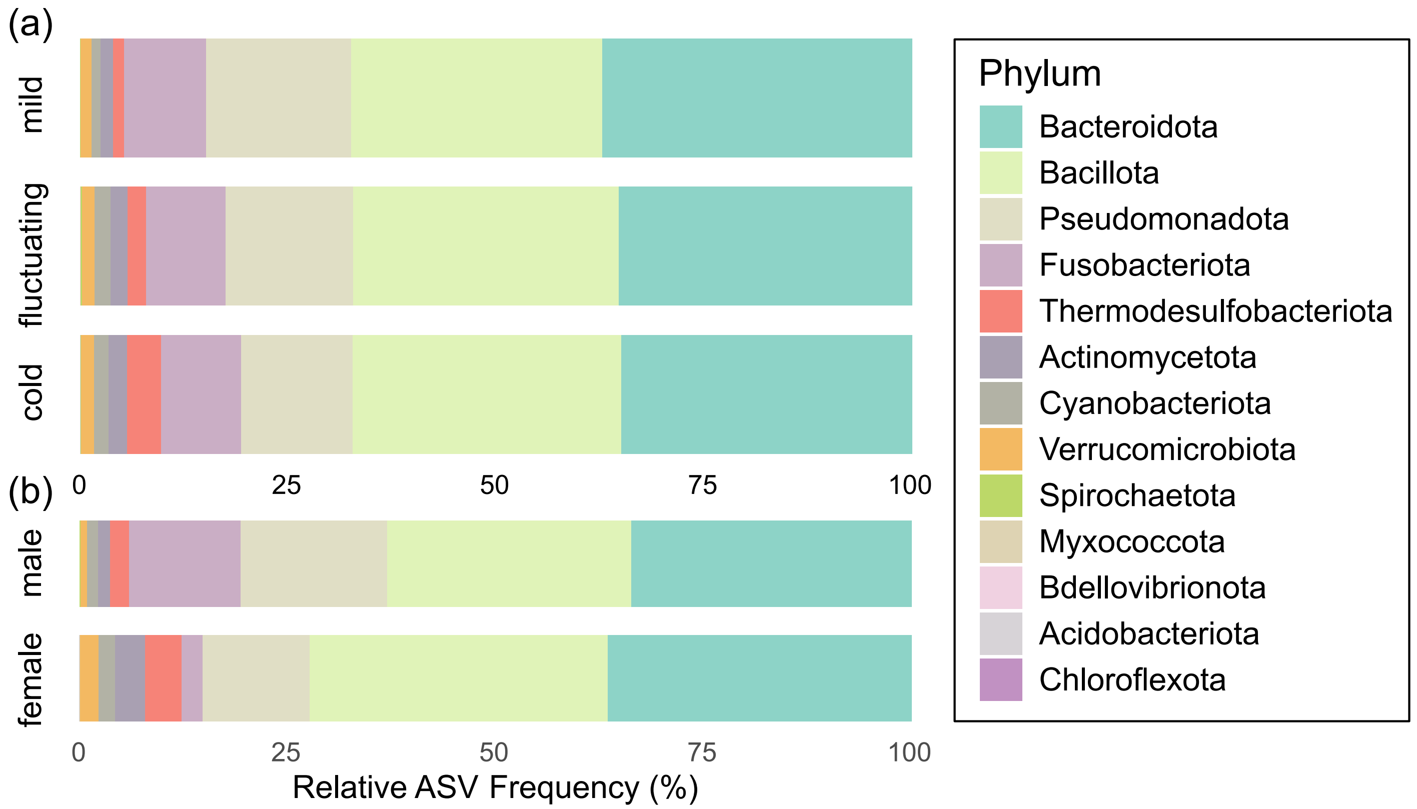

### Supplementary Figure S3

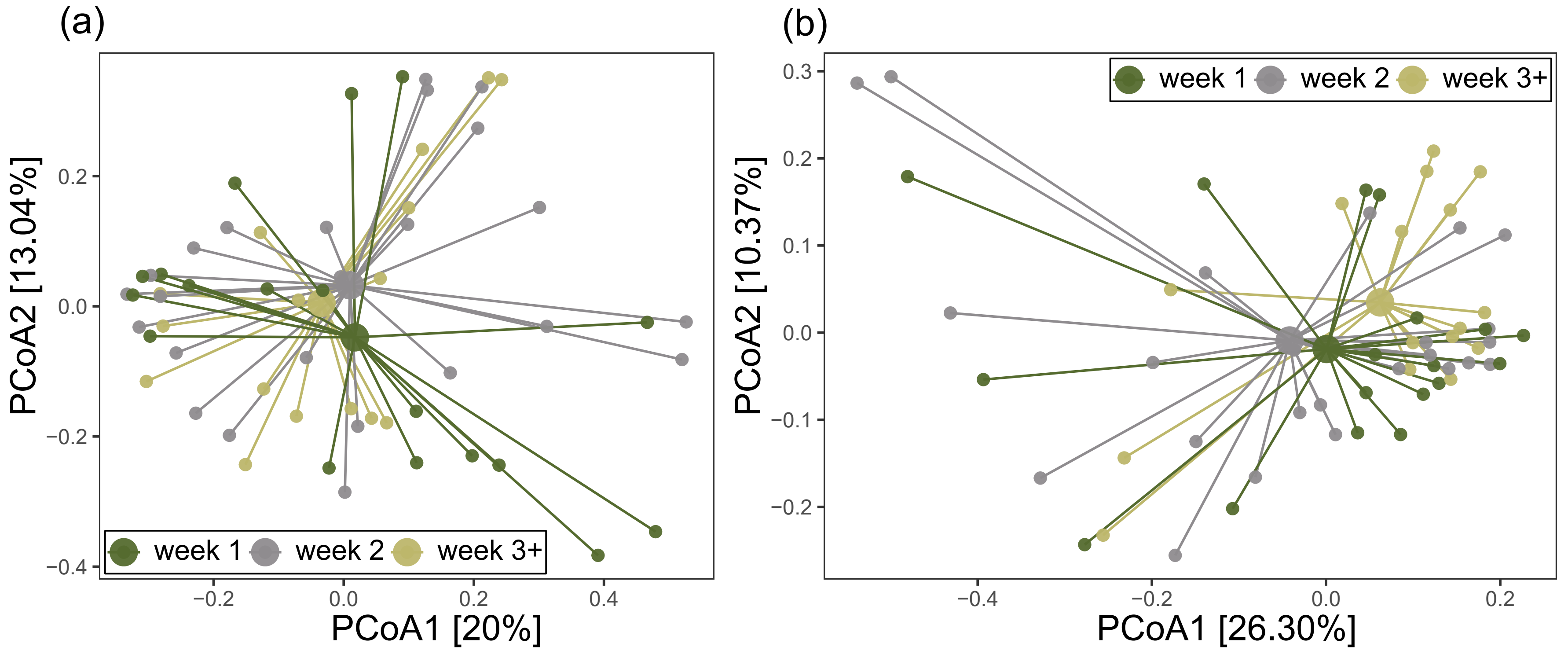

### Supplementary Figure S4

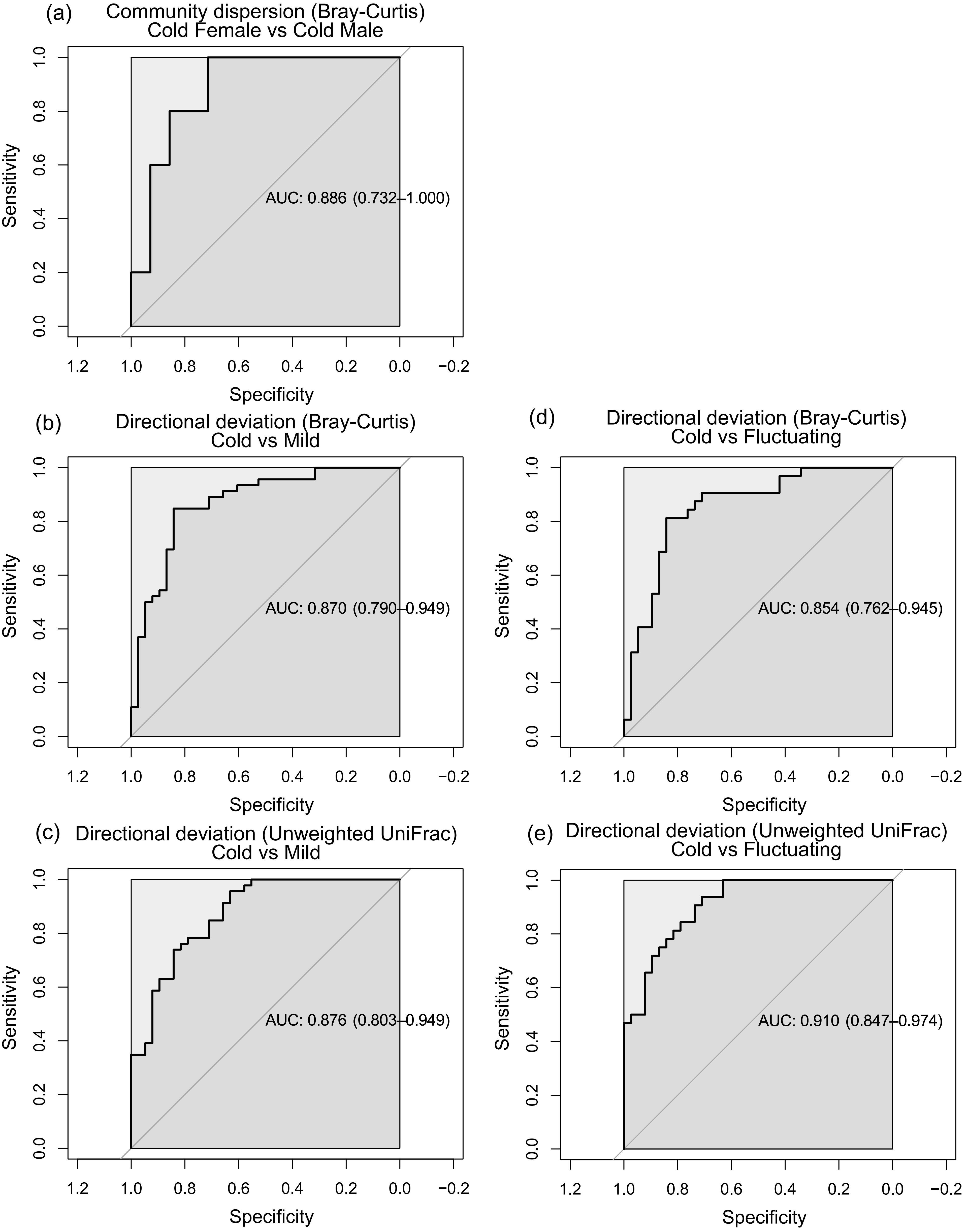

### Supplementary Figure S5

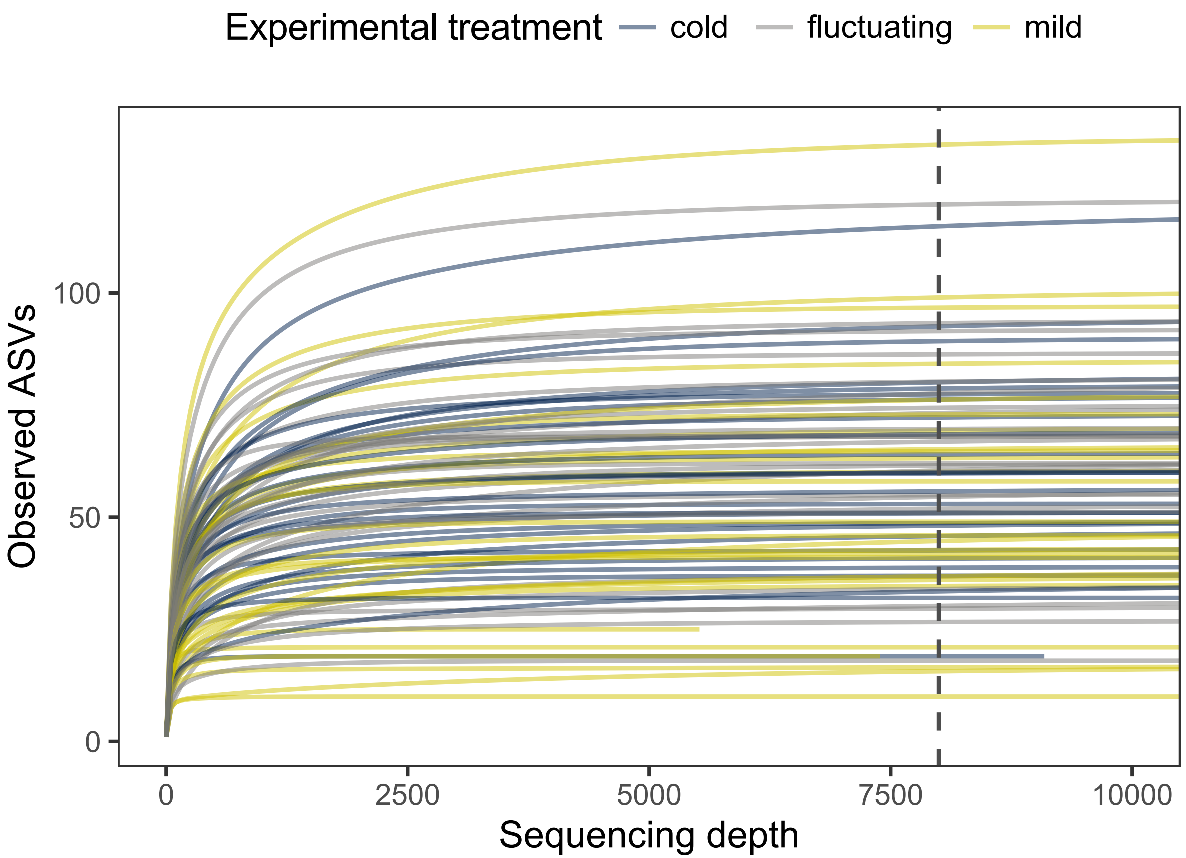
